# GEO2RNAseq: An easy-to-use R pipeline for complete pre-processing of RNA-seq data

**DOI:** 10.1101/771063

**Authors:** Bastian Seelbinder, Thomas Wolf, Steffen Priebe, Sylvie McNamara, Silvia Gerber, Reinhard Guthke, Jörg Linde

**Affiliations:** Research group Systems Biology / Bioinformatics, Leibniz Institute for Natural Product Research and Infection Biology – Hans-Knöll-Institute (HKI), Beutenbergstraße 11a, 07745 Jena, Germany; Research group PiDOMICS, Leibniz Institute for Natural Product Research and Infection Biology – Hans-Knöll-Institute (HKI), Beutenbergstraße 11a, 07745 Jena, Germany; Institute for Bacterial Infections and Zoonoses, Federal Research Institute for Animal Health – Friedrich-Loeffler-Institute, Naumburger Str 96a, 07743 Jena, Germany

**Keywords:** next-generation sequencing, RNA-seq, dual RNA-seq, differentially expressed genes, comparative transcriptomics, bioinformatics

## Abstract

In transcriptomics, the study of the total set of RNAs transcribed by the cell, RNA sequencing (RNA-seq) has become the standard tool for analysing gene expression. The primary goal is the detection of genes whose expression changes significantly between two or more conditions, either for a single species or for two or more interacting species at the same time (dual RNA-seq, triple RNA-seq and so forth). The analysis of RNA-seq can be simplified as many steps of the data pre-processing can be standardised in a pipeline.

In this publication we present the “GEO2RNAseq” pipeline for complete, quick and concurrent pre-processing of single, dual, and triple RNA-seq data. It covers all pre-processing steps starting from raw sequencing data to the analysis of differentially expressed genes, including various tables and figures to report intermediate and final results. Raw data may be provided in FASTQ format or can be downloaded automatically from the Gene Expression Omnibus repository. GEO2RNAseq strongly incorporates experimental as well as computational metadata. GEO2RNAseq is implemented in R, lightweight, easy to install via Conda and easy to use, but still very flexible through using modular programming and offering many extensions and alternative workflows.

GEO2RNAseq is publicly available at https://anaconda.org/xentrics/r-geo2rnaseq and https://bitbucket.org/thomas_wolf/geo2rnaseq/overview, including source code, installation instruction, and comprehensive package documentation.

## 1 INTRODUCTION

Organisms constantly change the expression of their genes to adapt to changes in the environment. With the help of transcriptomics scientists are able to study the gene expression of a given organism allowing to get insights in the interplay of genes dependent on environmental alterations. The primary goal is the detection of genes whose expression changes significantly between two or more conditions, and the function relationship of these genes. RNA sequencing (RNA-seq) offers a complete, fast, and cheap way to perform transcriptomics of single organisms using next-generation-sequencing technologies (Mardis, 2008). However, species do not exist in isolation. In fact, interspecies interactions are a major part of environmental adaptation. The special variant dual RNA-seq can be applied to analyse the transcriptome of two interacting species at the same time by separating their RNA *in silico* (Schulze et al., 2016; Wolf et al., 2018). The concept of dual RNA-seq can be further extended: triple RNA-seq allows investigating the interaction of three organisms, *e. g*. a host and two competing pathogens.

Both, the number of scientists applying RNA-seq and the number of published datasets have been growing exponentially ^1^ (Deelen et al., 2014). However, the bottlenecks in transcriptomics are the small number of experts able to pre-process and analyse RNA-seq data, and the small number of easy-to-use tools. A number of pipelines were published in R to handle this issue, but none of them includes the complete set of pre-processing steps and none exploits the available metadata fully.

Extensive utilisation of metadata is highly important. Wet-lab metadata, *e. g*. temperature or pH, and dry-lab metadata, *e. g*. p-value cut-offs or reference genome version, both influence the outcome of RNA –seq analysis and are often indispensable for the reproducibility of results (Rayner et al., 2006). Metadata help to decide if and how different datasets are comparable. Especially when many samples were used or different datasets were combined, metadata help to understand to keep track of the structure of a dataset. Knowledge of wet-lab metadata is essential for the interpretation computational results. To allow comparability of different analyses of the same datasets it is important to provide as much metadata as possible. Providing metadata for a possible subsequent analysis of the current data is as important as incorporating available metadata into a new analysis.

## 2 THE GEO2RNASEQ PIPELINE

GEO2RNAseq offers a complete analaysis of RNASeq data starting at FASTQ files or with a download from Gene Expression Omnibus (GEO).

At each step of the pipeline, the user has the possibility to change all parameters of the applied tools, which gives complete control over the processing steps. GEO2RNAseq is completely modular, allowing users to skip processing steps they do not need, integrate other R packages, or start the analysis at a later step. A restart-upon-error mechanism is implemented which detects incomplete and unprocessed samples. The mechanisms also allows the user to add new samples easily to the analysis without the need to re-run the most time consuming steps, i.e. quality control, mapping and gene abundance estimation. Every function is documented with a complete describing of input parameters, default values and output. Usually, only the paths to input files or The pipeline is highly scalable. Runnnig it with one thread/core on a laptop or on a server with multiple processors and threads/cores is a matter of changing one variable [`MAX_CPUS`] in the R pipeline script and results in heavy parallelisation of almost all processing steps. For cluster computing, the Rmpi package is installed with GEO2RNAseq and can be used to pre-process batches of read data on different machines. Figure 1 illustrates the workflow of GEO2RNAseq.

**Figure 1.**
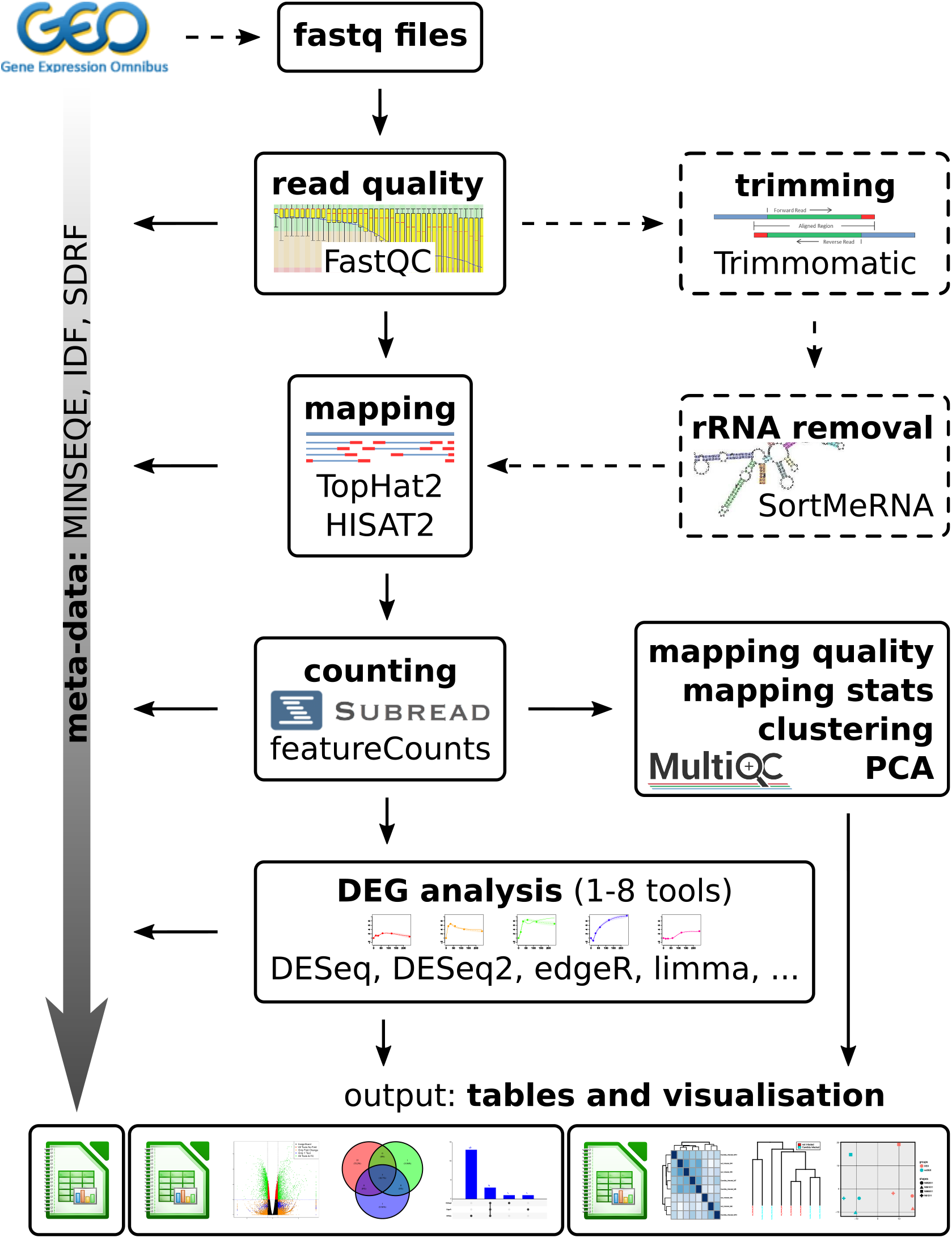
GEO2RNAseq workflow overview. The pipeline can download raw sequencing and metadata from the Gene Expression Omnibus (GEO) repository with a given GSE accession number. It automatically parses metadata according to the MINSEQE standard. It pre-processes raw sequencing data using many different tools. Some steps are optional, others offer alternative tools to apply. Differentially expressed genes are identified using up to eight different tools. By default, intermediate and final results are reported using tables in CSV and Excel format, various quality plots, hierarchical clustering and PCA, Venn diagrams, intersection bar plots, and volcano plots; more are available. Please note that this figure illustrates the *logical* order of processing steps as suggested by the authors. The arrows connecting each tool do not necessarily represent direct input/output interactions.

GEO2RNAseq automatically extracts metadata from GEO according to the MINSEQE standard hereby extending the IDF and SDRF formats (Rayner et al., 2006). If available from GEO, the following IDF features are retrieved automatically: investigation title, experiment class (*e. g*. RNA-seq, microarray), experimental description, experimental design according to the MGED ontology (*e. g*. “in_vitro_design”), experimental factors (*e. g*. strain or line), quality control information (*e. g*. “biological_replicate”), data release information, personal information of data submitter(s) and investigator(s) (*e. g*. Name, e-mail address, affiliation), and information about publications based on the dataset (*e. g*. PubMed ID). Again, if available from GEO, the following SDRF features are retrieved automatically per each sample: source information (*e. g*. organism part, sampling time point, culture conditions), applied protocols or standard operation procedures (SOP) for RNA extraction (*e. g*. extraction method), sequencing (*e. g*. technology type, read length, library source), and data processing (*e. g*. software parameters). Most importantly, the fold change assignment (“design matrix”), is extracted giving information on which comparisons of what samples have been or should be used for differentially expression analysis. The user can change or extend the metadata as necessary. During processing, the pipeline automatically extents the already available metadata by detailed information about each applied processing step and their parameters.

The pipeline starts with downloading raw sequencing data and corresponding metadata from GEO for a given GSE accession number. Next, it converts Sequence Read Archive (SRA) files into FASTQ files. Users can start directly with a set of FASTQ files as well. Quality check of FASTQ files is checked using FastQC^2^. Next, quality trimming of low-quality regions and adapter trimming is performed with Trimmomatic (Bolger et al., 2014). rRNA reads can be removed by using SortMeRNA (Kopylova et al., 2012) if required. SortMeRNA searches for rRNA sequences on a domain-of-life-basis, i.e. it distinguishes rRNA from bacterial, eukaryotic and archaeal origin. We implemented convenient wrappers to either (i) filter for rRNA from only one of the domains, (ii) filter for all three at the same time but keep a separate report, and (iii), to speed up SortMeRNA runtime, filter for all three as a whole and only report a total sum of rRNA content. At this point, users can decide to run FastQC again to check how trimming and rRNA removal improved the overall quality of the dataset. For mapping against a reference genome, the user can choose between TopHat2 (Kim et al., 2013) and HISAT2 (Kim et al., 2015). Other mappers, like STAR (Dobin et al., 2013) or BWA (Li and Durbin, 2009) can be easily integrated into the GEO2RNAseq workflow. In case of prokaryotic data, we suggest to use HISAT2 with the `--no-spliced-alignment` argument. To count the number of reads per gene, the pipeline uses featureCounts (Liao et al., 2013). As an indicator for rRNA contamination or high levels of globin-derived RNA in whole-blood datasets (Zhao et al., 2018), the pipeline automatically detects genes with particularly high read coverage. If such genes were detected, the user may consider to remove these genes from the count data, but this is not mandetory.

MultiQC (Ewels et al., 2016) is used to summarise quality control, read mapping and counting by creating interactive tables and various quality plots. GEO2RNAseq includes a MultiQC configuration file, which is optimised to work best with the pipeline’s output files and default directory structure. Next, comprehensive mapping statistics, including genome and transcriptome coverage, are calculated. Clustering of samples using different clustering methods (single-, complete-, average-linkage, median, centroid and all other methods as available in the `hclust` function) and visualizations (Figure 2 and Figure 3), and a principal component analysis (PCA, Figure 4) is performed and plotted.

Multiple different normalisation methods for count matrices (raw read counts), clustering, and PCA are implemented. These include counts per million (CPM), transcripts per million (TPM), read per kilobase million (RPKM), median by-ratio normalisation (MRN) (Anders and Huber, 2010), log-ratio, and variance stabilisation transformation (VST).

**Figure 2.**
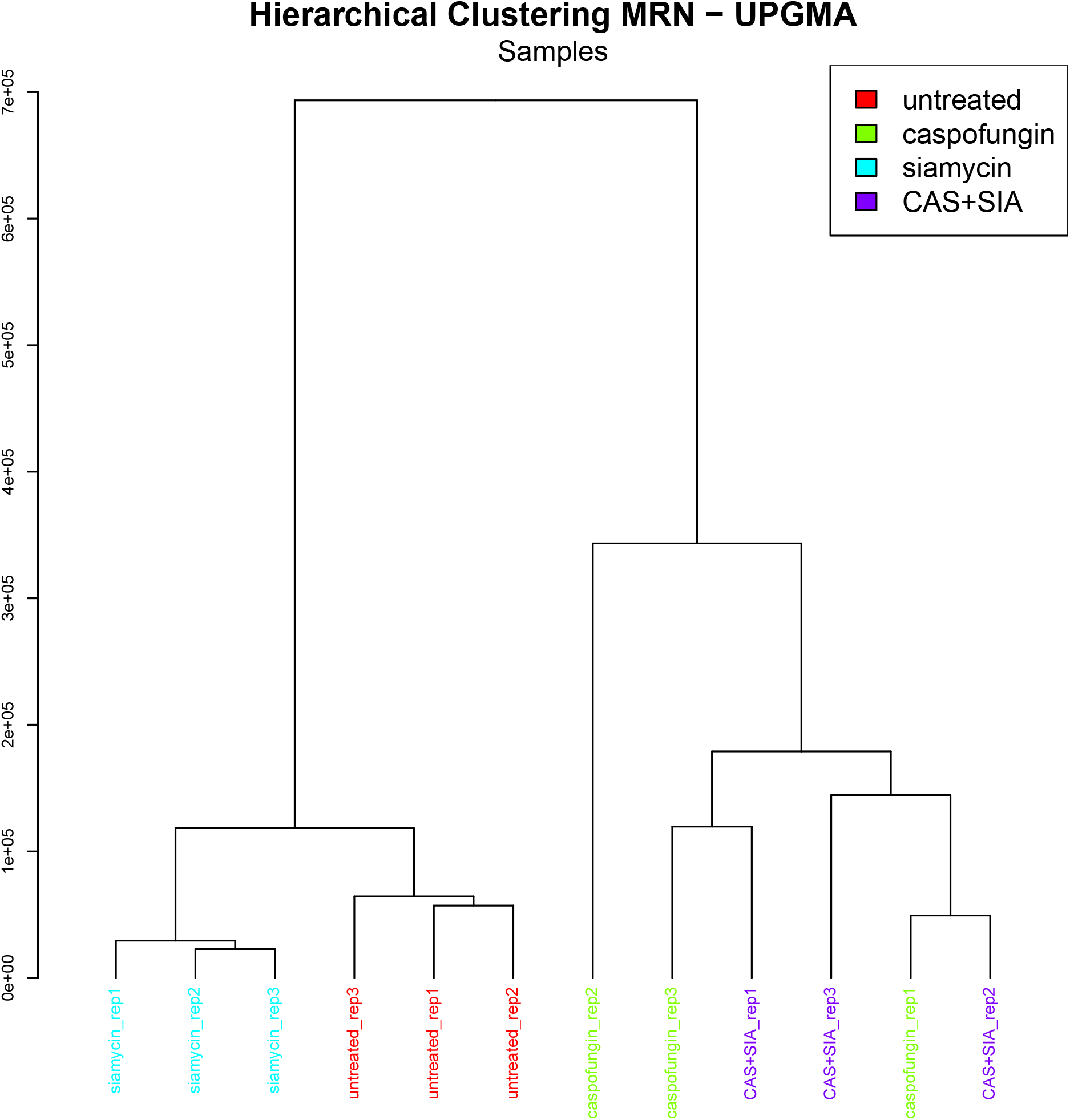
Hierarchical clustering of raw (not normalised) read count values. Samples can be coloured by user specified conditions or by conditions parsed from metadata. Here: Clear separation of samples treated with Caspofungin (green) or Caspofungin and Siamycin (CAS+SIA; purple) in the right subtree and samples treated with Siamycin only (blue) and the control sample (untreated; red) in the left subtree.

**Figure 3.**
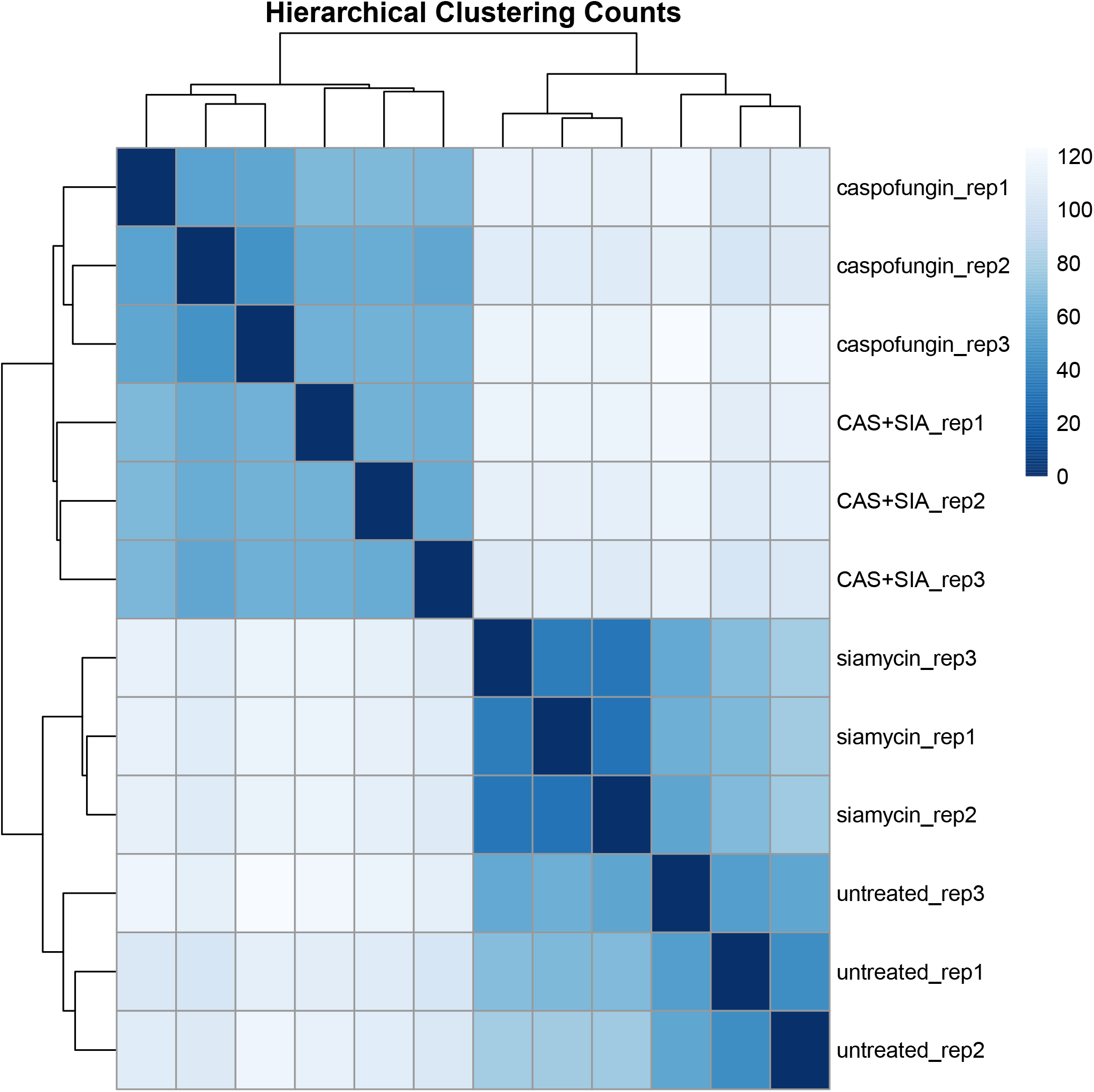
Similarity heatmap of MRN (Anders and Huber, 2010) normalised read count values with hierarchical clustering. The samples/replicates treated with Caspofungin only as well as Siamycin and Caspofungin (CAS+SIA) are in the upper left corner. The samples treated with Siamycin only and the control (untreated) are in the bottom right corner.

**Figure 4.**
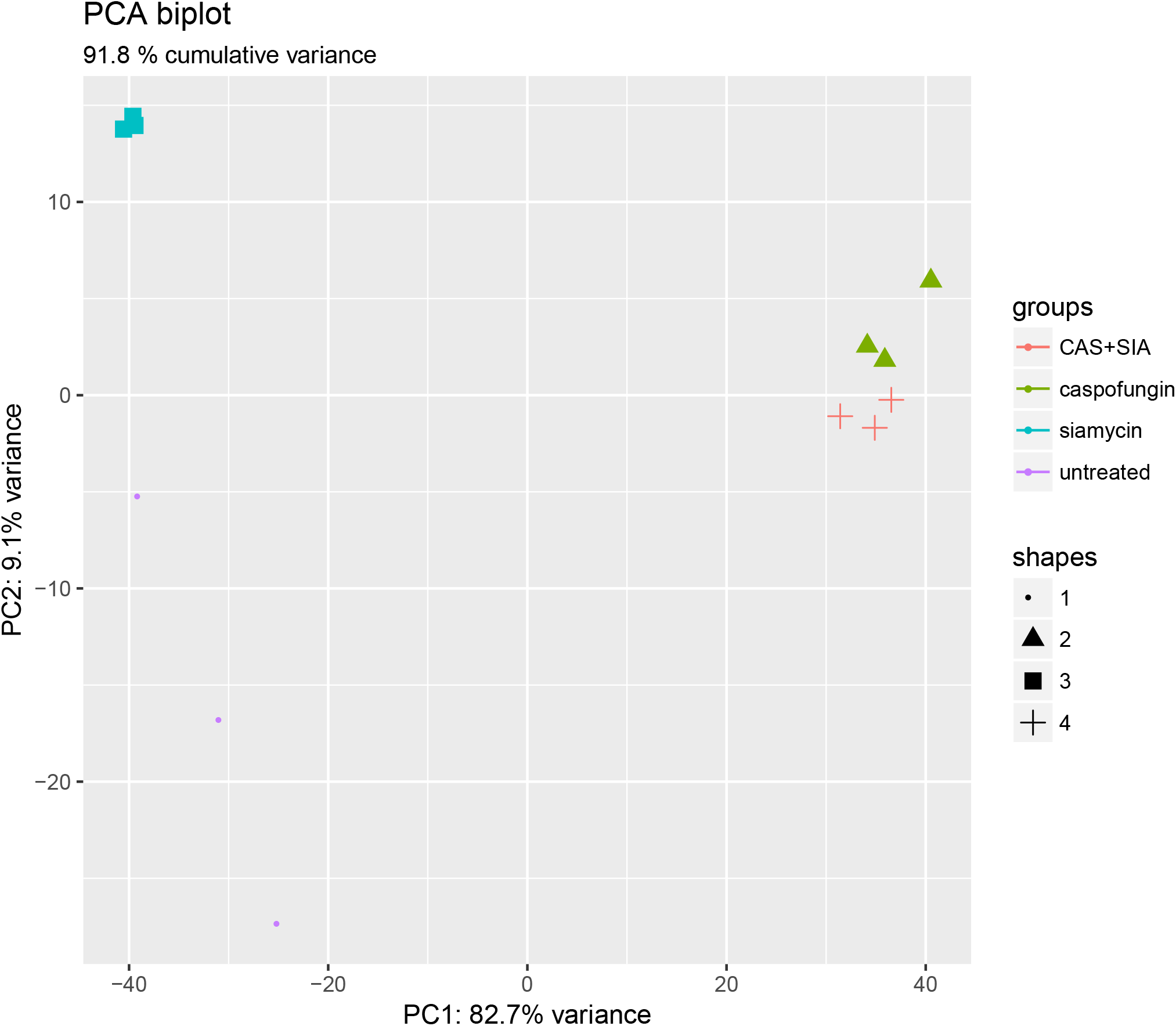
Principal component analysis (PCA) of MRN (Anders and Huber, 2010) normalised read count values showing a clear separation of samples treated with Caspofungin (green) or Caspofungin and Siamycin (CAS+SIA; red) on the right – and samples treated with Siamycin only (blue) and the control sample (untreated; purple) on the left. The three replicates of the control sample show a much higher variance, compared to the other samples. The plot can be further extended using ggplot2 in R.

Finally, GEO2RNAseq calculates significantly differentially expressed genes (DEGs) according to a given q-value cut-off and, optionally, log_2_ fold-change cut-off, for all comparisons defined manually by the user or automatically by using metadata. By default, the tools DESeq (Anders and Huber, 2012), DESeq2 (Love et al., 2014), edgeR (Robinson et al., 2010), and limma (McCarthy and Smyth, 2009) are execute concurrently and used to calculate adjusted p-values (q values) for each gene and comparison. In addition, baySeq (Hardcastle and Kelly, 2010), NOIseq (Tarazona et al., 2012), SAMseq (Tibshirani et al., 2011), and PoissonSeq (Li et al., 2012) are available. See Gierliński et al. (2015), Schurch et al. (2016), and Froussios et al. (2017) for in-depth analyses of strengths and weaknesses of various DEG tools. Moulos *et al*. Moulos and Hatzis (2015) demonstrate how the specificity, *i.e*. reducing type 2 errors for detected DEGs, is improved by using the maximum p-values across different statistical tools. We implemented the same strategy in GEO2RNAseq. Sensitivity can be increased by either using less stringent tools (e.g. only DESeq2 and edgeR) or increasing the q value threshold. Fold-change filtering can be applied on MRN, TPM or RPKM transformed abundance values as well.

The GEO2RNAseq vignette (see supplement file 1 `Vignette.html`) provides a comprehensive introduction to the basic concepts of an RNA-seq analysis and a step-by-step guide for using GEO2RNAseq. It explains the common steps of the RNA-seq pre-processing workflow and the different types of GEO datasets. Most importantly, the vignette describes each step of the GEO2RNAseq pipeline and the data structures introduced by this package. It includes the rationale behind each pre-processing step, a short description of each applied tool, example R codes, and the results produced by each code snippet.

Supplement file 2 `example_pipeline.R` may be used as a “minimal RNA-seq pipeline”. We also provide more comprehensive pipeline scripts for complete pre-processing of single (regular), dual, triple, and mixed single-end–paired-end RNA-seq datasets.

GEO2RNAseq combines 14 mandatory functions (see next section) that can be complemented with a collection of 132 functions for complete RNA-seq data pre-processing. GEO2RNAseq is implemented in R version 3.4.1 and publicly available at https://anaconda.org/xentrics/r-geo2rnaseq. The Conda^3^ package supports Linux and Mac systems. It includes a basic R installation as well as all R, external tool, and system dependencies required to run the pipeline. Therefore, it is very easy to install and use the pipeline within different computer environments. A tar ball for Conda-independent installation, source code, installation instructions, manual, and vignette are provided alongside the Conda package.

To ensure fast processing of RNA-seq data, GEO2RNAseq uses parallelisation, i. e. multi-threading and concurrent execution, wherever possible. The most time consuming and critical steps of RNA-seq data processing are handled by third-party software, e.g., Trimmomatic (Bolger et al., 2014), Sort-MeRNA (Kopylova et al., 2012), TopHat2 (Kim et al., 2013) or HISAT2 (Kim et al., 2015). Thus, run-time, accuracy and so on mainly depend on these third-party software products and are intensively discussed in their respective references. By parallelising the execution of third-party tools in addition to any intermediate R code, we are able to reduce the total runtime of the pipeline drastically. Therefore, the total execution time almost only depends on the size of the dataset, quality control and mapping of reads, the speed of the data drives, and the number of CPUs available to run GEO2RNAseq. The overall accuracy depends on the specifications of the third party software applied.

## 3 APPLICATION AND EXAMPLE RESULTS

To exemplarily present the capabilities of GEO2RNAseq, we re-analysed the RNA-seq dataset GSE55663 Valiante et al. (2015), which is freely available at GEO^4^, using only metadata provided within GSE55663. This dataset is based on RNA-seq results from of *Aspergillus fumigatus* treated with Caspofungin and Siamycin. It consists of three replicates for each condition, including a control condition. The R script for this re-analysis is provided in supplement `example_pipeline.R`. Furthermore, GEO2RNAseq has been successfully applied for the analysis of other single, dual, and also triple RNA-seq datasets.

For the complete pre-processing of GSE55663, we downloaded raw sequencing files and metadata using the function `getGEOdata()`. The automatically downloaded metadata can be found in supplement files 3 `GSE55663_SDRF.tsv` and 4 `GSE55663_IDF.tsv`. Next, we checked the sequencing quality of raw read data using the function `run_FastQC()`, performed adapter and quality trimming of the same read data using the function `run_Trimmomatic()`, and checked the quality of these trimmed reads. Trimmed reads were mapped to the reference genome of *A. fumigatus* strain A1163 using the function `run_Hisat2()`. The number of reads per gene were counted using `run_featureCounts()`. Reference genome and annotation files (in FASTA and GTF format) were downloaded from http://www.aspergillusgenome.org/. Only the download location of these files must be supplied to the functions. The index of the reference genome was created using `make_HiSat2_index()`. The latter required about 1 Gb of RAM. The results of all processing steps mentioned so far were summarised by the function `run_MultiQC()`. They can be found in supplement file `multiqc_report.zip`. GEO2RNAseq calculated detailed mapping statistics using the function `calc_mapping_stats()`, the result of which can be found in supplement file `mapping_stats.xls`. We then parsed the treatment information from the column ‘characteristics’. For normalisation, we used the function `get_mrn()`. After normalisation, we used `make_hclust_plot(), make_heat_clustering_plot()`, and `make_PCA_plot()` to generate the clustering (Figure 2, Figure 3) and PCA (Figure 4) plots. Based on the treatment conditions, we generated all pairwise tests using `create_design_matrix()`. Finally, we calculated differentially expressed genes for each test using `calculate_DEGs()` (Figure 5, Figure 6, Figure 7, Figure 8). For each comparison, this function creates Venn diagrams, intersection bar, vulcano and MA plots as well. In parallel, the pipeline collected metadata for each processing step, like tools, their respective versions and parameters. The resulting enriched metadata table is provided as supplement file `meta_data_final.csv`.

**Figure 5.**
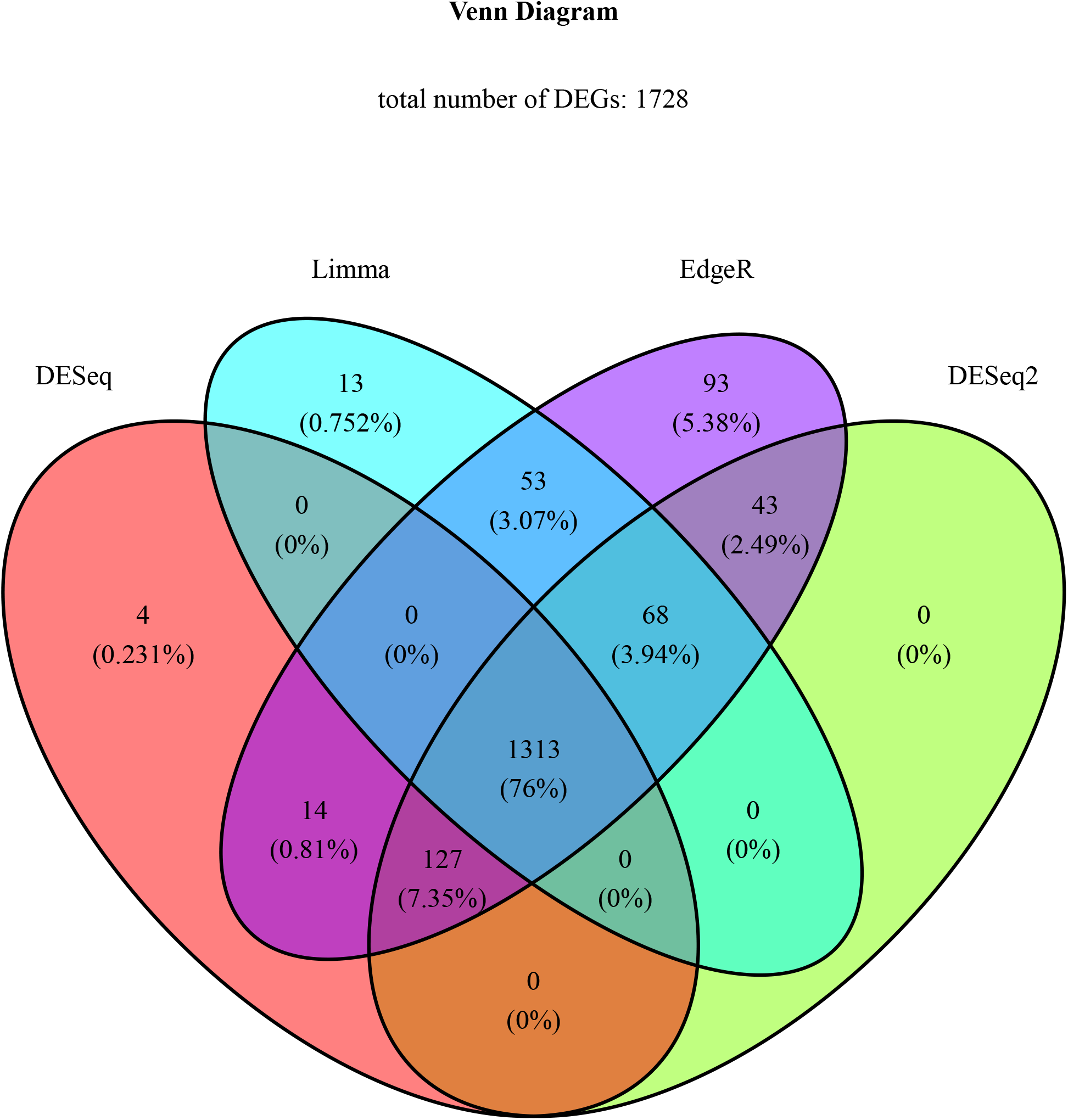
Venn diagram showing the overlap and difference of four sets of differentially expressed genes (DEGs) from the four DEG analysis tools DESeq (Anders and Huber, 2012), DESeq2 (Love et al., 2014), edgeR (Robinson et al., 2010), and limma (McCarthy and Smyth, 2009), which are used by default in GEO2RNAseq. Up to eight different DEG tools are supported. More than four are visualised using intersection bar plots (see Figure 6).

**Figure 6.**
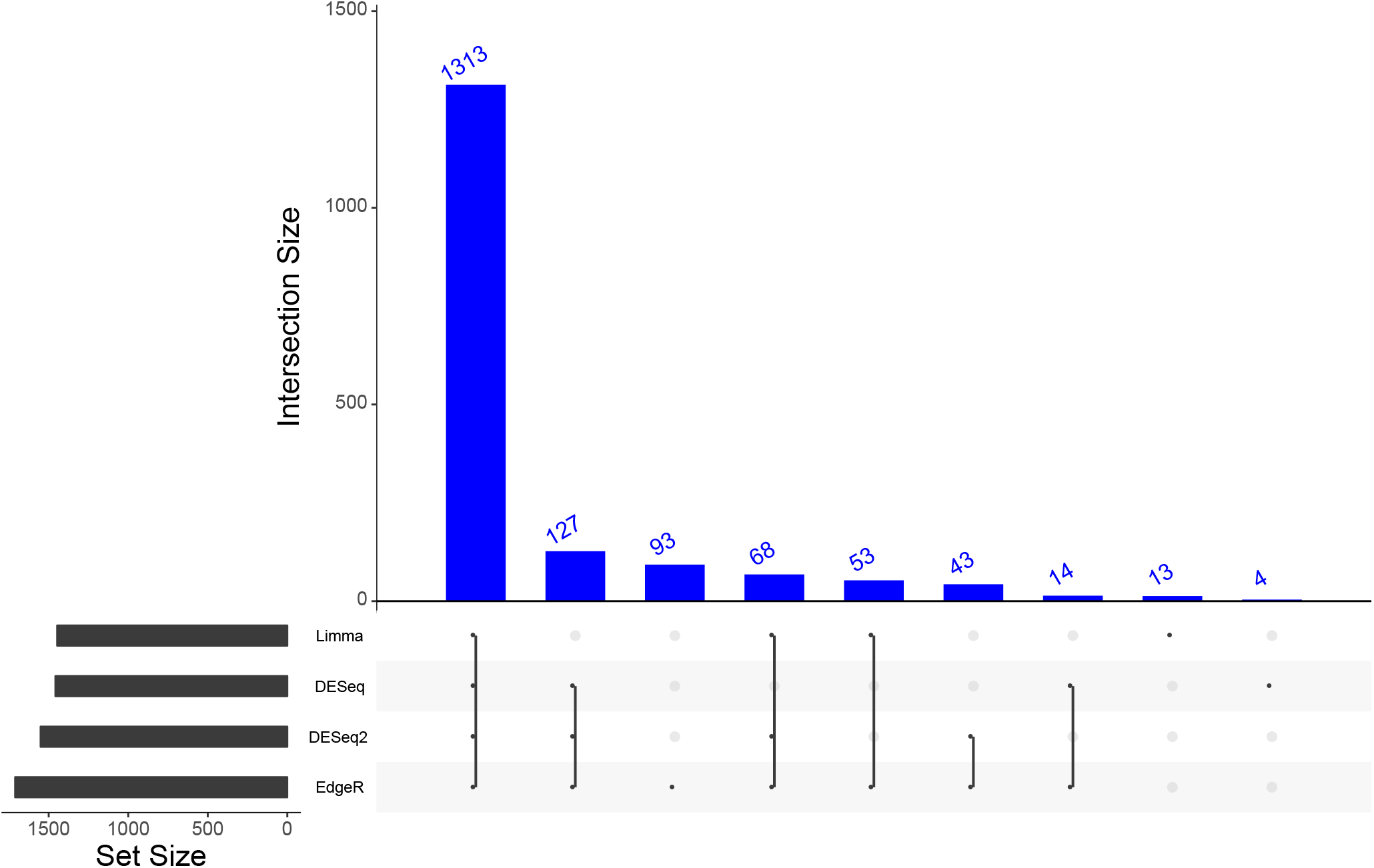
The intersection bar plot is an alternative for Venn diagrams and especially helpful if comparing more than four sets. Here: Four sets of differentially expressed genes (DEGs) from the four DEG analysis tools DESeq (Anders and Huber, 2012), DESeq2 (Love et al., 2014), edgeR (Robinson et al., 2010), and limma (McCarthy and Smyth, 2009), which are used by default in GEO2RNAseq. Up to eight different DEG tools are supported by GEO2RNAseq.

**Figure 7.**
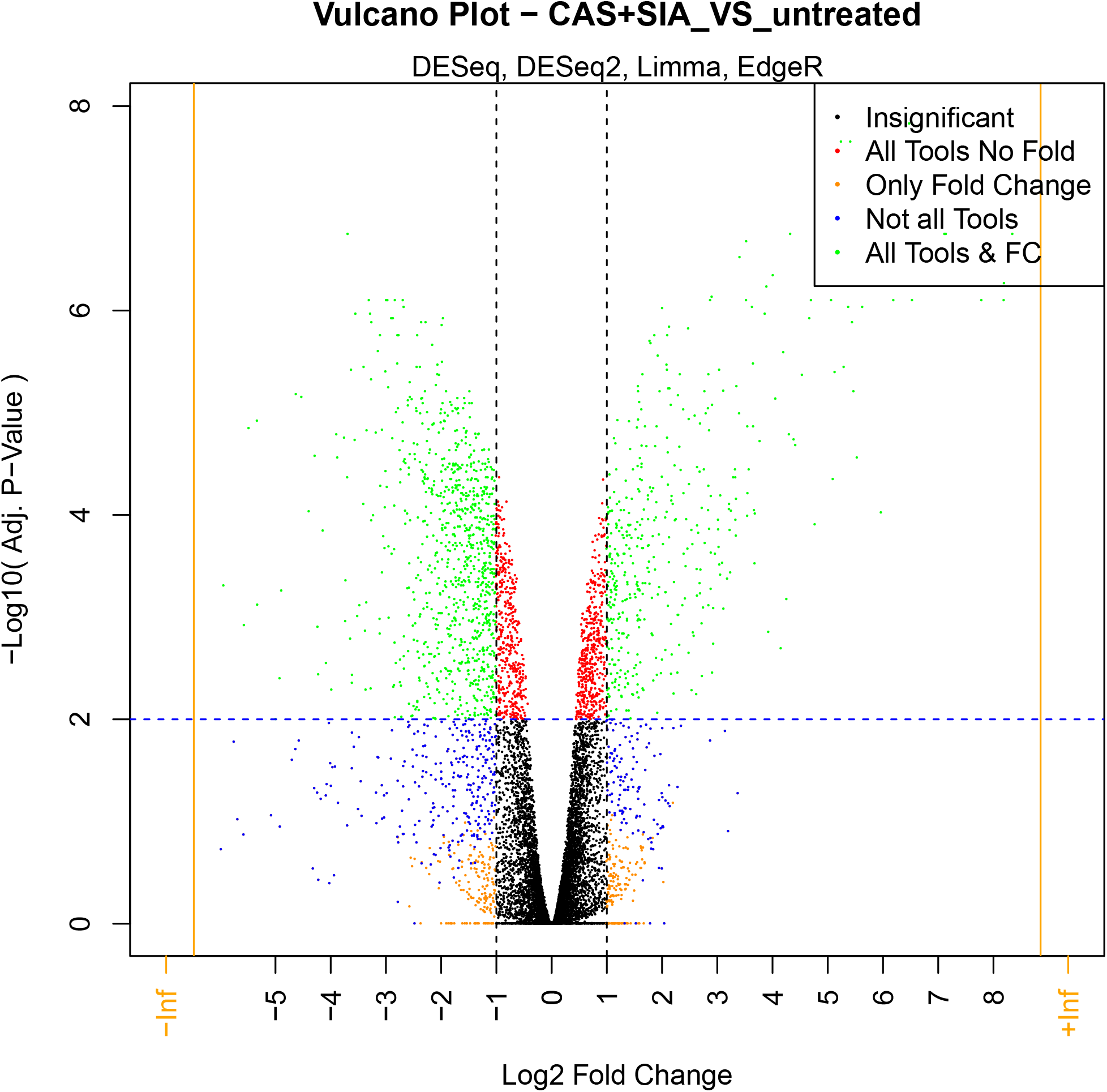
Volcano plot showing the distribution of log_2_ fold-changes and adjusted p-values from the DEG analysis of the comparison “control (untreated) against treatment with Caspofungin and Siamycin (CAS+SIA)”. When multiple DEG analysis tools are used, the maximum (“worst”) p-value per gene over these tools is used. Therefore, precision for DEGs is maximised and sensitivity minimised in the volcano plot. Genes with an adjusted p-value worse than the p-value cut-off (insignificant) are coloured black. If a fold-change filter was used during DEG analysis, vertical lines are added to indicate this threshold. Genes not achieving the fold-change threshold are coloured red. Insignificant genes achieving the fold-change cut-off are coloured orange. Blue coloured genes did not satisfy the adjusted p-value cut-off for all used tools, but may satisfy it for a subset of tools. Finally, genes satisfying both cut-offs for all tools are coloured green.

**Figure 8.**
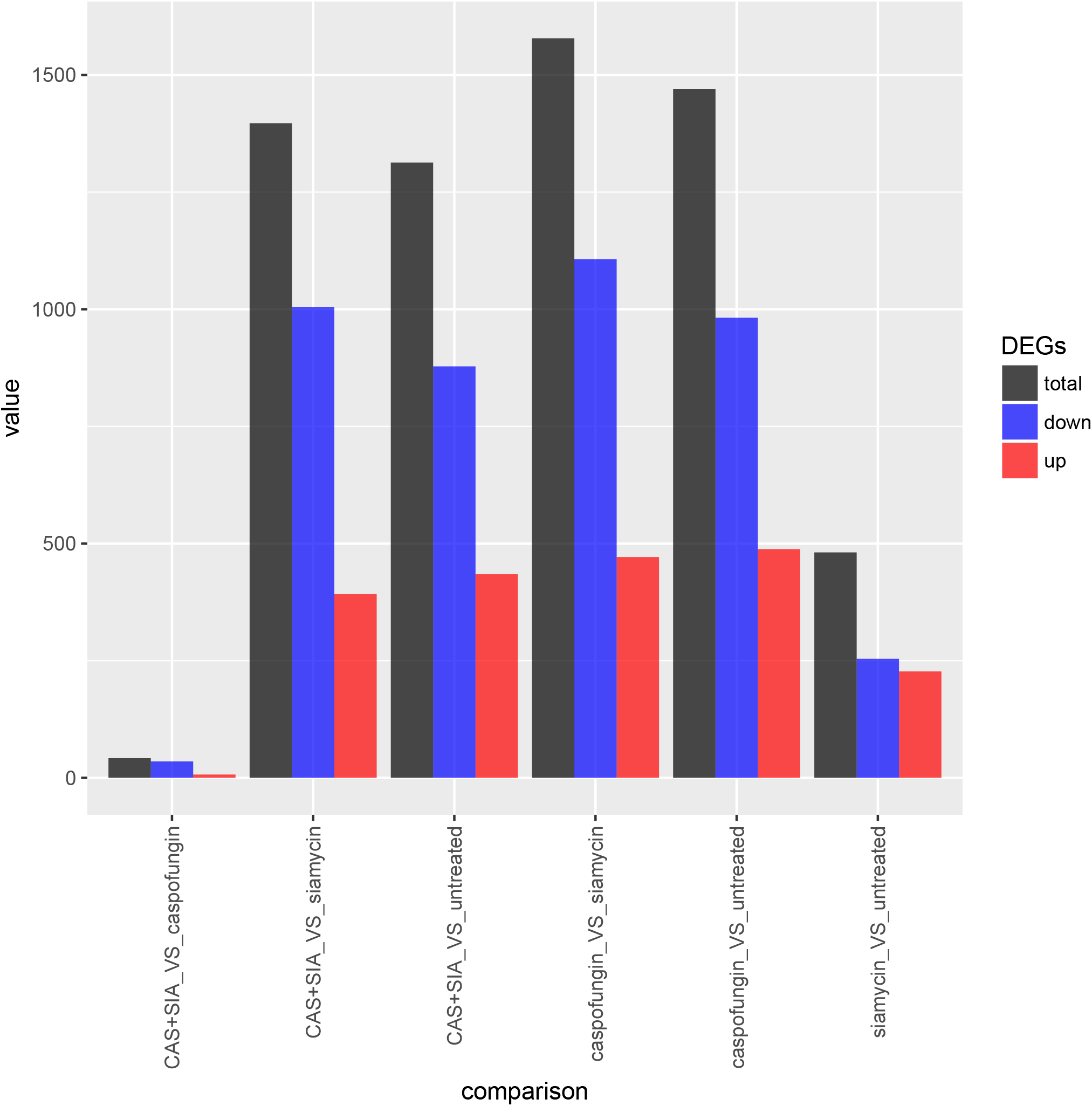
Bar plot showing the number of differentially expressed genes (DEGs) per comparison. This plot helps to quickly identify the subset of comparisons with a stronger treatment reaction according to the number of DEGs. It also shows weather a treatment induces more up- or downregulation. For the chosen comparisons, we see that a treatment with siamycin has a much weaker effect with respect to the number of DEGs when compared to caspofungin treatment. In addition, we see that the combination of caspofungin and siamycin treatment does not lead to substantially more DEGs. All treatments seem to induce an overall downregulation.

Intermediate and final results are saved in tabular form (Excel and CSV) and are supported by various plots (Table 1). The DEG analysis results include (i) normalised count values for each condition and for several normalisation methods, (ii) log fold-change values, and (iii) adjusted p-values from each applied DEG tool. More results are available as R objects and can be saved in the RData format (R workspace file) or can be further examined.

**Table 1.** Feature comparison of RNA-seq processing pipelines.

|  | GEO2RNAseq<sup>a</sup> | TRAPR<sup>b</sup> | EasyRNASeq<sup>c</sup> | READemption<sup>d</sup> |
| --- | --- | --- | --- | --- |
| wrapper language | R | R | R | Python |
| complete | yes | no | no | no<sup>e</sup> |
| input type | GEO<sup>f</sup>, FASTQ | counts | counts | FASTQ |
| quality control | FastQC | no | no | in-house |
| trimming | Trimmomatic | no | no | in-house |
| rRNA removal | SortMeRNA | no | no | no |
| mapping | TopHat2, HISAT2 | no | no | Segemehl |
| counting | featureCounts | no | bam2hits | in-house |
| counting result | raw, RPKM, TPM, MRN, rlog, vst | quantile, TMM, MRN | raw, RPKM, vst | raw, TNOAR<sup>g</sup> |
| mapping statistics | MultiQC, CSV | no | no | CSV |
| sample correlation | yes | yes | yes | yes |
| sample clustering | yes (Figure 2) | no | yes | no |
| sample heatmap | yes (Figure 3) | yes | yes | no |
| sample PCA | yes (Figure 4) | no | yes | no |
| DEG tools | 8<sup>h</sup> | 3 | 2 | 1 |
| DEG results |  |  |  |  |
| Venn diagram | yes (Figure 5) | no | no | no |
| UpSetR plot | yes (Figure 6) | no | no | no |
| Volcano plot | yes (Figure 7) | yes | no | yes |
| MA plot | yes | yes | no | yes |
| overview bar plot | yes (Figure 8) | no | no | no |
| modular | yes | yes | yes | no |
| dual/triple RNA-seq | yes | no | no | no |
| read/write metadata | yes | no | no | no |
| parallelisation | wrapper, external tools | no | yes | yes |
| Conda package | yes | no | yes | yes |

<sup>a</sup> https://anaconda.org/xentrics/r-geo2rnaseq<sup>b</sup> Lim et al. (2017)<sup>c</sup> Delhomme et al. (2012)<sup>d</sup> Förstner et al. (2014)<sup>e</sup> quality trimming of reads is incomplete<sup>f</sup> GSE accession<sup>g</sup> normalised by the total number of aligned reads (TNOR) and then multiplied by the lowest number of aligned reads of all considered libraries<sup>h</sup> four concurrent by default

GEO2RNAseq can be applied to analyse huge datasets, *e. g*. with more than 1000 samples, in a server environment or small datasets on a personal computer with limited capacities. We ran the example workflow (supplement `example_pipeline.R`), which consists of about 200 million raw reads in total, on a desktop computer. Intentionally we ran the computation on a 1 TB hard disk, which is much slower than a modern solid state drive. We allowed the pipeline to utilise up to six CPU cores. The entire system never allocated more than 8GB of memory. The total execution time was roughly 45 minutes, excluding the GEO download. Quality check with FastQC took 5 minutes, trimming with Trimmomatic 17 minutes,
 Mapping with HISAT2 about 20 minutes, featureCounts 2 minutes, PCA, clustering and statistical testing roughly 1 minute. This shows that the usage of GEO2RNAseq can be scaled up and applied to analyse vast datasets, *e. g*. with more than 1000 samples, in a server environment or small datasets on a personal computer with limited capacities.

Furthermore, GEO2RNAseq has been successfully applied for the analysis of other single, dual, and also triple RNA-seq datasets, for example by Schulze et al. (2015) and Sieber et al. (2018) (data not shown). For dual (and triple) RNA-seq, the pipeline requires one fitting reference genome in FASTA format and one annotation file in GTF format for each species. Annotation files must have the same attribute for gene IDs (usually the tag “gene_id”) or should be converted beforehand. Reference genomes are simply concatenated and then indexed together (using `make_HiSat2_index()`). Annotation files are concatenated together as well. The pipeline is executed the same as it was for single RNA-seq until gene abundance estimation was performed. The gene expression matrix can be separated into separate matrices for the two (or three) species. Hence, gene expression can then be analysed further per species or together. The latter, for example, is done when inferring inter-species gene expression correlations. In this case, normalization (e.g. MRN) is applied based using all genes from all species. This method removes the kind of bias that single RNA-seq of multiple species introduces by having different sequencing depths and hence normalization factors *between species*. In addition, the proportion of expression per species can be determined in dual RNA-seq which is not possible in single RNA-seq. Mapping statistics for dual RNA-seq are performed using `calc_dual_mapping_stats()` (or `calc_triple_mapping_stats()` for triple RNA-seq) which considers each species separately. Among other things, it calculates percentage contribution of RNA, exon and genome coverage per species. Example pipelines for performing dual and triple RNA-seq analyzes are supplied with the Geo2RNAseq R package.

## 4 COMPARISON TO OTHER RNA-SEQ PIPELINES

The very popular Galaxy workflow engine (Afgan et al., 2016) offers a scientific workflow, data integration, and analysis platform. It primarily aims to make computational biology accessible to scientists without programming skills. Today, Galaxy is a well-established platform, but installation, adaptation, and augmentation is still challenging. To achieve simplicity in usage, flexibility and output options of tools are limited when compared to their full capabilities. Using non-implemented tools and performing other follow up analyses requires to manually integrate and import such tools into Galaxy and to manually export the results from it. This difficulty potentially causes a barrier for efficient and error-free data processing when using novel tools.

The R software environment^5^ offers an easy framework for performing simple and complex statistics. Users have access to a huge number of packages for transcriptome data analysis, data interpretation, plotting and more. In this sense, R is much more flexible than Galaxy, but requires at least some programming skills of the user. In fact, many important steps of RNA-seq data processing, like the detection of differentially expresses genes (DEGs), are natively implemented in R.

Two pipelines for processing of RNAseq data exist and and were widely used. The Total RNA-Seq Analysis Package for R (TRAPR) (Lim et al., 2017) is a partial RNA-seq pipeline implemented in R. However, does not include mapping and counting, and uses counts per genes as input. Given counts, TRAPR performs all following steps necessary for the detection of DEGs, but the statistical analysis is limited to DESeq2 and edgeR.

EasyRNASeq (Delhomme et al., 2012) is a package available through Bioconductor^6^. Again, it can only be used after initial mapping was performed. As well the statistical analysis is limited to DESeq2 and edgeR.

In computer science, the programming language Python is often used in conjunction or as an alternative to R. READemption (Förstner et al., 2014) is a Python pipeline for processing differential RNA-seq data. However, quality trimming of reads is incomplete, it uses only DESeq2 for statistical analysis, and the processing steps are fixed. READemption utilised Segemehl (?) for mapping, which is rarely used as of today.

None of the R or Python pipelines mentioned before are able to use raw sequencing data available in the Gene Expression Omnibus (GEO) repository. None of them handles metadata according to the MINSEQE^7^ standard, and none of them can handle dual or triple RNA-seq datasets. See Table 1 for a detailed feature comparison.

In conclusion, GEO2RNAseq is complete and modular RNAseq pre-processing pipeline. In contrast to the other pipelines mentioned, users can process data in standardised ways or fine-tune the processing steps using all capabilities that R offers. Between each step, some of which are optional, the user may choose to use other R code or packages to fit their particular needs. GEO2RNAseq ensures this by keeping input and output of processed data comprehensive, but simple. Finally, GEO2RNAseq is highly scalable because it makes heavy use of parallelisation in form of concurrent execution and multi-threading.

## Supporting information

Comprehensive Code Vignette

Minimal Pipeline Example

## AUTHOR CONTRIBUTIONS

JL designed the project, workflow, and supported acquisition and analysis of data. BS, TW, JL, SP, SM, and SG programmed and shipped the tool. BS and TW wrote the manuscript. JL and RG contributed on writing the manuscript. All authors read, revised and approved the final manuscript.

## ACKNOWLEDGEMENTS

This work was supported by the Deutsche Forschungsgemeinschaft (DFG) CRC/Transregio 124 “Pathogenic fungi and their human host: Networks of interaction”, subproject INF (JL, TW), as well as the Free State of Thuringia and the European Social Fund 2016 FGR 0053 (JL, BS). We thank Maximilian Collatz and Peter Großmann for testing GEO2RNAseq and Iris Lüke for critically revising the language of the manuscript.

## Footnotes

1 https://www.ncbi.nlm.nih.gov/sra/docs/sragrowth/

2 https://www.bioinformatics.babraham.ac.uk/projects/fastqc

3 (Ana-) Conda is a package, dependency, and environment management systemfor many programming languages and operating systems (https://conda.io/docs/)

4 https://www.ncbi.nlm.nih.gov/geo/query/acc.cgi?acc=GSE55663

5 https://www.r-project.org

6 https://bioconductor.org

7 http://fged.org/projects/minseqe

## REFERENCES

Afgan, E., Baker, D., van den Beek, M., Blankenberg, D., Bouvier, D., Čech, M., et al. (2016). The Galaxy platform for accessible, reproducible and collaborative biomedical analyses: 2016 update. Nucleic Acids Research 44, W3–W10. doi:10.1093/nar/gkw343

Anders, S. and Huber, W. (2010). Differential expression analysis for sequence count data. Genome biology 11, R106

Anders, S. and Huber, W. (2012). Differential expression of rna-seq data at the gene level–the deseq package. Heidelberg, Germany: European Molecular Biology Laboratory (EMBL)

Bolger, A. M., Lohse, M., and Usadel, B. (2014). Trimmomatic: a flexible trimmer for Illumina sequence data. Bioinformatics 30, 2114–2120. doi:10.1093/bioinformatics/btu170

Deelen, P., V Zhernakova, D., de Haan, M., van der Sijde, M., Bonder, M. J., Karjalainen, J., et al. (2014). Calling genotypes from public rna-sequencing data enables identification of genetic variants that affect gene-expression levels

Delhomme, N., Padioleau, I., Furlong, E. E., and Steinmetz, L. M. (2012). easyrnaseq: a bioconductor package for processing rna-seq data. Bioinformatics 28, 2532–2533

Dobin, A., Davis, C. A., Schlesinger, F., Drenkow, J., Zaleski, C., Jha, S., et al. (2013). Star: ultrafast universal rna-seq aligner. Bioinformatics 29, 15–21

Ewels, P., Magnusson, M., Lundin, S., and Käller, M. (2016). MultiQC: summarize analysis results for multiple tools and samples in a single report. Bioinformatics 32, 3047–3048. doi:10.1093/bioinformatics/btw354

Förstner, K. U., Vogel, J., and Sharma, C. M. (2014). READemption – a tool for the computational analysis of deep-sequencing-based transcriptome data. Bioinformatics 30, 3421–3423. doi:10.1093/bioinformatics/btu533

Froussios, K., Schurch, N. J., Mackinnon, K., Gierlinski, M., Duc, C., Simpson, G. G., et al. (2017). How well do RNA-Seq differential gene expression tools perform in a eukaryote with a complex transcriptome? bioRxiv, 090753 doi:10.1101/090753

Gierliński, M., Cole, C., Schofield, P., Schurch, N. J., Sherstnev, A., Singh, V., et al. (2015). Statistical models for RNA-seq data derived from a two-condition 48-replicate experiment. Bioinformatics 31, 3625–3630. doi:10.1093/bioinformatics/btv425

Hardcastle, T. J. and Kelly, K. A. (2010). bayseq: empirical bayesian methods for identifying differential expression in sequence count data. BMC bioinformatics 11, 422

Kim, D., Langmead, B., and Salzberg, S. L. (2015). HISAT: a fast spliced aligner with low memory requirements. Nature Methods 12, 357–360. doi:10.1038/nmeth.3317

Kim, D., Pertea, G., Trapnell, C., Pimentel, H., Kelley, R., and Salzberg, S. L. (2013). TopHat2: accurate alignment of transcriptomes in the presence of insertions, deletions and gene fusions. Genome Biology 14, R36. doi:10.1186/gb-2013-14-4-r36

Kopylova, E., Noé, L., and Touzet, H. (2012). SortMeRNA: fast and accurate filtering of ribosomal RNAs in metatranscriptomic data. Bioinformatics 28, 3211–3217. doi:10.1093/bioinformatics/bts611

Li, H. and Durbin, R. (2009). Fast and accurate short read alignment with burrows–wheeler transform. Bioinformatics 25, 1754–1760

Li, J., Witten, D. M., Johnstone, I. M., and Tibshirani, R. (2012). Normalization, testing, and false discovery rate estimation for rna-sequencing data. Biostatistics 13, 523–538

Liao, Y., Smyth, G. K., and Shi, W. (2013). The subread aligner: fast, accurate and scalable read mapping by seed-and-vote. Nucleic Acids Research 41, e108

Lim, J. H., Lee, S. Y., and Kim, J. H. (2017). TRAPR: R Package for Statistical Analysis and Visualization of RNA-Seq Data. Genomics & Informatics 15, 51–53. doi:10.5808/GI.2017.15.1.51

Love, M. I., Huber, W., and Anders, S. (2014). Moderated estimation of fold change and dispersion for rna-seq data with deseq2. Genome biology 15, 550

Mardis, E. R. (2008). Next-Generation DNA Sequencing Methods. Annual Review of Genomics and Human Genetics 9, 387–402. doi:10.1146/annurev.genom.9.081307.164359

McCarthy, D. J. and Smyth, G. K. (2009). Testing significance relative to a fold-change threshold is a TREAT. Bioinformatics 25, 765–771. doi:10.1093/bioinformatics/btp053

Moulos, P. and Hatzis, P. (2015). Systematic integration of RNA-Seq statistical algorithms for accurate detection of differential gene expression patterns. Nucleic Acids Research 43, e25–e25. doi:10.1093/nar/gku1273

Rayner, T. F., Rocca-Serra, P., Spellman, P. T., Causton, H. C., Farne, A., Holloway, E., et al. (2006). A simple spreadsheet-based, MIAME-supportive format for microarray data: MAGE-TAB. BMC Bioinformatics 7, 489. doi:10.1186/1471-2105-7-489

Robinson, M. D., McCarthy, D. J., and Smyth, G. K. (2010). edger: a bioconductor package for differential expression analysis of digital gene expression data. Bioinformatics 26, 139–140

Schulze, S., Henkel, S. G., Driesch, D., Guthke, R., and Linde, J. (2015). Computational prediction of molecular pathogen-host interactions based on dual transcriptome data. Frontiers in Microbiology 6. doi:10.3389/fmicb.2015.00065

Schulze, S., Schleicher, J., Guthke, R., and Linde, J. (2016). How to Predict Molecular Interactions between Species? Frontiers in Microbiology 7. doi:10.3389/fmicb.2016.00442

Schurch, N. J., Schofield, P., Gierliński, M., Cole, C., Sherstnev, A., Singh, V., et al. (2016). How many biological replicates are needed in an RNA-seq experiment and which differential expression tool should you use? RNA 22, 839–851. doi:10.1261/rna.053959.115

Sieber, P., Voigt, K., Kämmer, P., Brunke, S., Schuster, S., and Linde, J. (2018). Comparative Study on Alternative Splicing in Human Fungal Pathogens Suggests Its Involvement During Host Invasion. Frontiers in Microbiology 9. doi:10.3389/fmicb.2018.02313

Tarazona, S., García, F., Ferrer, A., Dopazo, J., and Conesa, A. (2012). Noiseq: a rna-seq differential expression method robust for sequencing depth biases. EMBnet. journal 17, pp- 18

Tibshirani, R., Chu, G., Narasimhan, B., and Li, J. (2011). samr: SAM: Significance Analysis of Microarrays. R package version 2.0

Valiante, V., Monteiro, M. C., Martín, J., Altwasser, R., Aouad, N. E., González, I., et al. (2015). Hitting the Caspofungin Salvage Pathway of Human-Pathogenic Fungi with the Novel Lasso Peptide Humidimycin (MDN-0010). Antimicrobial Agents and Chemotherapy 59, 5145–5153. doi:10.1128/AAC.00683-15

Wolf, T., Kämmer, P., Brunke, S., and Linde, J. (2018). Two’s company: studying interspecies relationships with dual RNA-seq. Current Opinion in Microbiology 42, 7–12. doi:10.1016/j.mib.2017.09.001

Zhao, S., Zhang, Y., Gamini, R., Zhang, B., and Schack, D. (2018). Evaluation of two main rna-seq approaches for gene quantification in clinical rna sequencing: polya+ selection versus rrna depletion. Scientific reports 8, 4781

