## Supplementary material for "GEO2RNAseq: An easy-to-use R pipeline for complete pre-processing of RNA-seq data": Comprehensive Code Vignette

#### *2019-04-03*

###### Package

Geo2RNAseq 0.99.13

### Contents

- 1 Introduction
  - 1.1 GEO overview
    - 1.1.1 Platforms
    - 1.1.2 Samples
    - 1.1.3 Series
    - 1.1.4 Datasets
  - 1.2 Acquiring and Setting Up Data
  - 1.3 Access package data
  - 1.4 List of Tools Found by the Package
  - 1.5 Acquire Raw Data from GEO
  - 1.6 Metadata
- 2 Quality Check of Raw Read Data
- 3 Trimming
- 4 Quality Check of Filtered Read Data
- 5 Mapping
  - 5.1 Indexing
  - 5.2 Read Alignments
- 6 Gene Abundance Estimation
  - 6.1 Annotation
  - 6.2 Counting
- 7 Mapping Statistics
  - 7.1 Explanation of Mapping Stats Table
- 8 MultiQC
- 9 Clustering and PCA
  - 9.1 Load Count Data
  - 9.2 Hierarchical Clustering
  - 9.3 Correlation
  - 9.4 Principle Component Analysis
- 10 Design Matrix
- 11 DEG Analysis
- 12 Updating Metadata
- 13 Archive Results
- 14 Example Pre-processing Workflows (Pipelines)
- 15 Session Info

### 1 Introduction

RNA-sequencing (RNA-seq) using next-generation sequencing (NGS) technologies has become a standard technology for analyzing gene expression profiles. Finding significantly, differentially expressed genes between two or more groups of interest is a common goal of transcriptomic research. RNA-seq allows to perform such analysis on a genome-wide scale.

RNA-seq experiments often focus on the analysis of gene expression changes within *one* organism. It is also possible to sequence *two* interacting species at the same time, e.g. during host-pathogen interaction. *GEO2RNAseq* works equally well with “ordinary” RNA-seq datasets and so called “dual RNA-seq” datasets. Only for metagenomics, where *thousands* of different species are sequenced at the same time, we suggest to look for another pipeline.

RNA-seq raw data is given in the form of short sequencing “reads” in FASTQ format. For experiments available in the Gene Expression Omnibus (GEO) repository, FASTQ files can be downloaded automatically using functions provided by *GEO2RNAseq*. It will also download experimental metadata from GEO, such as sample processing information, sequencing platform, and so on.

Given a set of FASTQ files or “samples”, base-quality and adapter trimming is performed to improve downstream analysis. High-quality reads are then aligned against a reference genome sequence. This process is called “read mapping”" or simply “mapping”. An appropriate reference genome must be selected to match the organism from which the samples were created. Given the location of each mapped read, gene abundances or gene expression levels are estimated based on known gene locations. Gene locations are annotated using standardized formats, such as GTF, and are available for most reference genomes. The abundance of each gene is estimated for each sample. At last, samples and their gene abundances are grouped together based on experimental conditions (e.g., treatment and control). Statistical testing is performed to evaluate abundance changes of genes between those two groups, resulting in one or more lists of (significantly) differentially expressed genes.

In summary, GEO2RNAseq offers the following features:

- automatic download from GEO
- handling of metadata before and during pre-processing
- quality control with *FastQC*
- trimming with *Trimmomatic*
- mapping with *TopHat2* or *HISAT2*
- read counting with *featureCounts* (from *Rsubread*)
- combination of reports and log files using *MultiQC*
- detection of DEGs with any combination of *DESeq*, *DESeq2*, *edgeR*, *limma*, *NOISeq*, *baySeq*, *PoissonSeq* or SAMseq (from *samr*)

In *GEO2RNAseq*, these steps can be performed with little manual labour. It uses a collection of tools (see this section) and manages experimental and bioinformatics metadata (temperature, pH, tool calls, tool versions, …). Each processing step is performed by a single function call and returns basic data objects such as `data.frames` and `list` objects. This allows the user to make use of other R packages and processing steps. In general, the pipeline can be started and stopped at any step if the neccessary input data is already present.

*GEO2RNAseq* makes use of parallelisation wherever possible to achieve a maximal speed of pre-processing.

The user can also use other implemented functions to easily calculate mapping statistics (e.g. the number of reads mapped to exons) or perform clustering and PCA on count data using different normalization methods (e.g. TPM, RPKM, vst).

Different statistical tools may calculate different significance values (p-values) for the same gene. Intersections of gene lists from the different DEG tools are visualized using Venn Diagrams and Intersection Bar plots (from *UpSetR*).

#### 1.1 GEO overview

Gene Expression Omnibus (GEO) is a public repository that archives and distributes microarray and high-throughput sequencing data. This includes metadata, e.g. information about sample preparation, project goals and findings, sequencing machines, per-sample conditions, and so on. GEO data records are divided into 4 main protocols: GPL (platforms), GSM (samples), GSE (series), GDS (datasets). *GEO2RNAseq* only supports data and metadata download from GSE data records. These are used to summarize experiments and group sample information together.

##### 1.1.1 Platforms

A platform record contains a description of the array or sequencer. Each platform record is assigned a unique and stable GEO accession number (*GPLxxx*). A platform may reference many samples. Platforms are indexed and searchable using the Entrez GEO datasets interface. GPL accessions cannot be used with this package.

##### 1.1.2 Samples

A sample record contains a description of how the sample was obtained and processed. A data table with normalized abundance measurements for each feature on the corresponding platform is usually included, as well as links to corresponding raw data files. Each sample record is assigned a unique and stable GEO accession number (*GSMxxx*). Samples are usually part of a GEO series and are used for acquiring raw read data and the corresponding metadata.

##### 1.1.3 Series

A series record defines a set of related samples (GSM) and how these are related. A series provides a description of the experiment as a whole. Each series record is assigned a unique and stable GEO accession number (*GSExxx*). These are the main accession numbers to be used with *GEO2RNAseq*. The GSM accessions corresponding to a GSE entry are accessed, raw sample data is downloaded, and all acquired information is summarized in form of a simple table.

##### 1.1.4 Datasets

GEO datasets (*GDSxxx*) are curated sets of GEO sample data. A GDS record represents a collection of biologically and statistically comparable GEO samples. Samples within a GDS refer to the same Platform, that is, they share a common set of probe elements.
*To this data (2018-27-02), there are no curated NGS datasets. Therefore, GDS accessions are not supported. This may change in the future.*

#### 1.2 Acquiring and Setting Up Data

We demonstrate the main features of *GEO2RNAseq* using data packages from Bioconductor. Because raw data is usually large, we use *two different datasets* throughout this vignette. Pre-processing, including counting, is performed on a few thousend reads generated from the *Drosophila melanogaster* genome 3 chromosome 4 (BDGP Release 5, dm3, UCSC, April 2006). The reads and reference chromosome are included in this package. The corresponding genome was acquired from the *pasillaBamSubset* package.

We perform hierarchical clustering, correlation and principal component analysis (PCA), as well as differential gene expression analysis, using the count matrix from the *airway* package. The use and access of GEO accessions will be demonstrated using the corresponding accession *GSE52778*.

#### 1.3 Access package data

```
library("Geo2RNAseq")
```

The FASTQ files and metadata files are supplied inside the *extdata* directory of this package. All output files are saved to the following directory:

```
pkgDir <- system.file("extdata", package = "Geo2RNAseq")
# all files created in this vignette will be written to this directory
outDir <- file.path(pkgDir, "outData")
outDir
```

```
## [1] "/home/basti/.conda/envs/r-dev/lib/R/library/Geo2RNAseq/extdata/outData"
```

#### 1.4 List of Tools Found by the Package

Before using *GEO2RNAseq*, make sure that external programs listed in the table below are installed and can be found from within the R session. We tested *GEO2RNAseq* on the stated versions but the pipeline can easily be adapted to updates. The following code displays all tools, including their binaries and versions. In addition, it shows system variables which can be changed outside the R environment. On UNIX systems, this refers to exporting a variable from the “.bashrc” (or similar) file. If the system variable for a tool is defined, it should point to the corresponding binary file. In that case, standard search paths are ignored. Otherwise, the default paths for executables (e.g., */usr/bin/* or *PATH* on UNIX) are searched.

Table 1: List of external tools

|  | version | system.variable |
| --- | --- | --- |
| bowtie2 | 2.2.2 | BOWTIE2\_EXEC |
| fastq-dump | 2.8.2 | FASTQ\_DUMP\_EXEC |
| fastqc | 0.11 | FASTQC\_EXEC |
| hisat2 | 2.1 | HISAT2\_EXEC |
| multiqc | 1.5 | MULTIQC\_EXEC |
| samtools | 0.1.19 | SAMTOOLS\_EXEC |
| sortmerna | 2.1 | SORTMERNA\_EXEC |
| trimmomatic | 0.36 | TRIMMOMATIC\_EXEC |
| tophat2 | 2.1.0 | TOPHAT2\_EXEC |

```
list_executables()
```

```
## 
## === List of installed tools ===
## --> fastq-dump
##     sys var: FASTQ_DUMP_EXEC = 
##     exec: '/home/basti/.conda/envs/r-dev/bin/fastq-dump.2.8.2'
##     version: fastq-dump : 2.8.2
## --> multiqc
##     sys var: MULTIQC_EXEC = 
##     exec: '/home/basti/.conda/envs/r-dev/bin/multiqc'
##     version: 1.5
## --> trimmomatic
##     sys var: TRIMMOMATIC_EXEC = 
##     exec: '/home/basti/.conda/envs/r-dev/share/trimmomatic-0.36-5/trimmomatic'
##     version: 0.36
## --> samtools
##     sys var: SAMTOOLS_EXEC = 
##     exec: '/home/basti/.conda/envs/r-dev/bin/samtools'
##     version: 1.8 (using htslib 1.8)
## --> sortmerna
##     sys var: SORTMERNA_EXEC = 
##     exec: '/home/basti/.conda/envs/r-dev/bin/sortmerna'
##     version:   SortMeRNA version 2.1b, 03/03/2016
## --> bowtie2
##     sys var: BOWTIE2_EXEC = 
##     exec: '/home/basti/.conda/envs/r-dev/bin/bowtie2'
##     version: 2.2.8
## --> tophat2
##     sys var: TOPHAT2_PATH = 
##     exec: '/home/basti/.conda/envs/r-dev/bin/tophat2'
##     version: TopHat v2.1.1
## --> hisat2
##     sys var: HISAT2_EXEC = 
##     exec: '/home/basti/.conda/envs/r-dev/bin/hisat2'
##     version: 2.1.0
## --> fastqc
##     sys var: FASTQC_EXEC = 
##     exec: '/home/basti/.conda/envs/r-dev/opt/fastqc-0.11.7/fastqc'
##     version: 0.11.7
```

#### 1.5 Acquire Raw Data from GEO

As mentioned above, GEO series (GSE accessions) records are used. These contain not only the samples (and their metadata) but also general information about the experiment. The function ‘getGEOdata’ accesses all GSM entries corresponding to the given GSE, downloads the raw data in form of SRA files, converts these to FASTQ files using fastq-dump and combines all acquired information into three metadata tables. The returned list has three keywords. “META” refers to metadata in MINSEQE format, “SDRF” and “IDF” are the same metadata but filtered and changed to fit the corresponding standard.
*In the following example, the metadata is used as packaged with GEO2RNAseq. In general, you would call the function ‘getGEOdata’:*

```
# With network access and time, you would access GEO like this:
# geo_dat <- getGEOdata(accession = "GSE52778", outDir = file.path(outDir, "GSE52778"))
# This will create 3 table-like files, download 16 SRA files and convert them to
# 16 FASTQ files.
geo_dat <- getGeoDemoDat(pkgDir)
print(names(geo_dat))
```

```
##  [1] "SDRFfile"  "IDFfile"   "METAfile"  "SDRF"      "META"      "sra_files"
##  [7] "se_files"  "pe_files"  "se_index"  "pe_index"
```

```
# You may check the content like this:
# View(geo_dat$META)
# View(geo_dat$SDRF)

# the files can also be loaded using standard functions or functions beginning with 'parse_'
META <- read.csv(geo_dat$METAfile, header = TRUE, sep = "\t", as.is = TRUE)
SDRF <- parse_SDRF(geo_dat$SDRFfile)
```

```
## Fold Change Assignment column not found. Ignored.
```

```
## Warning in parse_SDRF(geo_dat$SDRFfile): Did not find any design matrix
## information. You have to add this manually to the SDRF file.
```

#### 1.6 Metadata

Metadata is stored in form of `data.frame`s. For compatibility, *GEO2RNAseq* works best with the MINSEQE and SDRF formats.

An empty, minimal metadata file with 3 rows can be created using:

```
make_meta_table(outDir = outDir, nrRows = 3, overwrite = TRUE, as.xls = TRUE)
```

```
## Create empty metadata table
```

```
## $file
## [1] "/home/basti/.conda/envs/r-dev/lib/R/library/Geo2RNAseq/extdata/outData/meta_data.xls"
## 
## $table
##   Specimen Organism Strain Date Library.Strategy  Dual Data.File
## 1     FILL     FILL     NA   NA           SINGLE FALSE     FILL1
## 2     FILL     FILL     NA   NA           SINGLE FALSE     FILL2
## 3     FILL     FILL     NA   NA           SINGLE FALSE     FILL3
```

```
# or as CSV
# make_meta_table(outDir = outDir, nrRows = 3, overwrite = TRUE, as.xls = FALSE)
```

The user may add more information to the resulting Excel/CSV file. Afterwards, it can be read in again:

```
parsed_meta_info <- parse_meta_xls(file.path(outDir, "meta_data.xls"))
```

```
## Warning in parse_meta_xls(file.path(outDir, "meta_data.xls")): Did not find
## any design matrix information. You have to add this manually to the XLS file.
```

```
# or for CSV
# parsed_meta_info <- parse_meta_csv(file.path(outDir, "meta_data.csv"))
```

This code throws a warning because the parser attempts to read design matrix information, which is not yet available, but will be discussed in a later section.

In this section, the metadata object is updated based on the following processing steps.

### 2 Quality Check of Raw Read Data

Let’s start with the actual data pre-processing: the quality check of raw read data. Here, we use the artificial reads. They are shipped with this packaged. Reads were generated from the Drosphila reference sequence mentioned in the introduction. To run *FastQC*, only the paths to the FASTQ files must be supplied to its wrapper function:

```
fq <- system.file("extdata", "synthetic.fastq.gz", package = "Geo2RNAseq")
fq
```

```
## [1] "/home/basti/.conda/envs/r-dev/lib/R/library/Geo2RNAseq/extdata/synthetic.fastq.gz"
```

```
rawQualDir <- file.path(outDir, "quality", "raw")
rawQualDir
```

```
## [1] "/home/basti/.conda/envs/r-dev/lib/R/library/Geo2RNAseq/extdata/outData/quality/raw"
```

```
fq_raw_res <- run_FastQC(files = fq, outDir = rawQualDir, cpus = 2, workers = 2)
fq_raw_res
```

```
## $calls
## [1] "/home/basti/.conda/envs/r-dev/opt/fastqc-0.11.7/fastqc /home/basti/.conda/envs/r-dev/lib/R/library/Geo2RNAseq/extdata/synthetic.fastq.gz --outdir /home/basti/.conda/envs/r-dev/lib/R/library/Geo2RNAseq/extdata/outData/quality/raw --threads 2"
## 
## $version
## [1] "v0.11.7"
## 
## $tool
## [1] "fastqc"
```

You may check the HTML report(s) in the output directory before proceeding.

### 3 Trimming

Trimming includes the removal of adapter sequences and the removal of low quality sequences. For this step, *GEO2RNAseq* utilizes the tool *Trimmomatic*. By default, leading and trailing low quality bases of a read are removed (base-quality below 4). In addition, window trimming is performed (by default: window size 15 bp, average quality 25, from both ends). This means that the average quality of a sliding window is checked. The sequence is cut if the quality falls below a given cut-off. Finally, reads shorter than a given length (30 nucleotides) are removed completely.

See `?run_Trimmomatic` for all parameters that can be changed. For most cases, `is.paired`, `qualcut`, and `adapters` may be set. Set `is.paired` to `TRUE` when working with paired-end data and `FALSE` otherwise. `qualcut` defines the minimum quality during the window size trimming. If the average quality is below this value, the entire scanned sequence is cut off. `adapters` is the path to a FASTA file containing the adapter sequences used during the sequencing process. By default, the function uses the ILLUMINA PE-3 adapter sequences. Adapter files/sequences are shipped with *Trimmomatic* and with this package.

```
trimDir  <- file.path(outDir, "fastq")
trim_res <- run_Trimmomatic(fq, outDir = trimDir, is.paired = FALSE, compress = TRUE, cpus = 2, workers = 2)
```

```
## Trimming ...
```

```
## /home/basti/.conda/envs/r-dev/share/trimmomatic-0.36-5/trimmomatic SE -threads 2 -phred33 /home/basti/.conda/envs/r-dev/lib/R/library/Geo2RNAseq/extdata/synthetic.fastq.gz /home/basti/.conda/envs/r-dev/lib/R/library/Geo2RNAseq/extdata/outData/fastq/synthetic.trimo.fq.gz ILLUMINACLIP:/home/basti/.conda/envs/r-dev/share/trimmomatic-0.36-5/adapters/TruSeq3-SE.fa:2:30:10 LEADING:3 TRAILING:3 SLIDINGWINDOW:15:25 MINLEN:30 2>&1
```

```
trim_res
```

```
## $files
## [1] "/home/basti/.conda/envs/r-dev/lib/R/library/Geo2RNAseq/extdata/outData/fastq/synthetic.trimo.fq.gz"
## 
## $calls
## [1] "/home/basti/.conda/envs/r-dev/share/trimmomatic-0.36-5/trimmomatic SE -threads 2 -phred33 /home/basti/.conda/envs/r-dev/lib/R/library/Geo2RNAseq/extdata/synthetic.fastq.gz /home/basti/.conda/envs/r-dev/lib/R/library/Geo2RNAseq/extdata/outData/fastq/synthetic.trimo.fq.gz ILLUMINACLIP:/home/basti/.conda/envs/r-dev/share/trimmomatic-0.36-5/adapters/TruSeq3-SE.fa:2:30:10 LEADING:3 TRAILING:3 SLIDINGWINDOW:15:25 MINLEN:30 2>&1"
## 
## $input
## [1] 1000
## 
## $surviving
## [1] 1000
## 
## $log
## [1] "/home/basti/.conda/envs/r-dev/lib/R/library/Geo2RNAseq/extdata/outData/fastq/synthetic.trimlog"
## 
## $tool
## [1] "trimmomatic"
```

### 4 Quality Check of Filtered Read Data

It is advisable to check the impact of the trimming step on the quality of the read data. Run *FastQC* again, with `extend = TRUE`. This changes the output name so that the tool MultiQC (used at a later step) can differentiate reports of raw and, e.g., trimmed data. Otherwise, the reports will be combined.

```
fq <- trim_res$files
fq
```

```
## [1] "/home/basti/.conda/envs/r-dev/lib/R/library/Geo2RNAseq/extdata/outData/fastq/synthetic.trimo.fq.gz"
```

```
filterQualDir <- file.path(outDir, "quality", "filter")
fq_raw_res <- run_FastQC(files = fq, outDir = filterQualDir, extend = TRUE, cpus = 2, workers = 2)
```

Again, you may check the HTML report(s) in the output directory before proceeding.

### 5 Mapping

The process of aligning reads to a reference sequence, e.g. a genome, is called “mapping”. *GEO2RNAseq* allows to map reads using either TopHat2 or HISAT2. *HISAT2* is much faster than *TopHat2* and uses less memory, usually with little or no loss in sensitivity. *For your own data set, you must first acquire a reference genome FASTA file fitting your organism of interest. Such references can usually be found at, e.g., NCBI Genbank, Ensembl, JGI, or CGD. In the following, we continue with chromosome 4 of the Drosophila reference genome dm3*.

#### 5.1 Indexing

The genome sequence must be indexed first. Depending on which mapping tool you want to apply, *TopHat2* or *HISAT2*, run one of the following wrapper functions. The index files are created in the same directory as the genome. In this example, we use only chromsome 4 of the Drosophila reference genome dm3.

```
genomeFile <- system.file("extdata", "dm3_chr4.fa", package = "Geo2RNAseq")
genome <- file.path(outDir, "genome", basename(genomeFile))
dir.create(file.path(outDir, "genome"), recursive = TRUE)
file.copy(from = genomeFile, to = genome)
```

```
## [1] TRUE
```

```
# make index for TopHat2
tophat_index <- make_Tophat_index(genomeFile = genome)
```

```
## Make bowtie2 index using:
## /home/basti/.conda/envs/r-dev/bin/bowtie2-build
```

```
tophat_index
```

```
## $outpref
## [1] "/home/basti/.conda/envs/r-dev/lib/R/library/Geo2RNAseq/extdata/outData/genome/dm3_chr4"
## 
## $call
## [1] "/home/basti/.conda/envs/r-dev/bin/bowtie2-build /home/basti/.conda/envs/r-dev/lib/R/library/Geo2RNAseq/extdata/outData/genome/dm3_chr4.fa /home/basti/.conda/envs/r-dev/lib/R/library/Geo2RNAseq/extdata/outData/genome/dm3_chr4"
```

```
# OR: make index for HISAT2
hisat_index <- make_HiSat2_index(genomeFile = genome)
```

```
## Make HISAT2 index using:
## /home/basti/.conda/envs/r-dev/bin/hisat2-build
```

```
hisat_index
```

```
## $outpref
## [1] "/home/basti/.conda/envs/r-dev/lib/R/library/Geo2RNAseq/extdata/outData/genome/dm3_chr4"
## 
## $call
## [1] "/home/basti/.conda/envs/r-dev/bin/hisat2-build -p 1 /home/basti/.conda/envs/r-dev/lib/R/library/Geo2RNAseq/extdata/outData/genome/dm3_chr4.fa /home/basti/.conda/envs/r-dev/lib/R/library/Geo2RNAseq/extdata/outData/genome/dm3_chr4"
```

The path to index files are given to mapping tools without file extension. The function `tools::file_path_sans_ext()` removes file extensions.

```
map_index <- tools::file_path_sans_ext(genome)
map_index
```

```
## [1] "/home/basti/.conda/envs/r-dev/lib/R/library/Geo2RNAseq/extdata/outData/genome/dm3_chr4"
```

#### 5.2 Read Alignments

For the actual mapping, choose between TopHat2 and HISAT2. HISAT version 2.10.0 or higher is advised. See `?run_Tophat` and `?run_Hisat2` for additional parameters. In most cases, `is.paired` and `anno` may be set. Set `is.paired` to `TRUE` when working with paired-end data and `FALSE` otherwise. `anno` is only used for *TopHat2*. Supply the path to a annotation file, i.e. a GFF or GTF formatted file describing the transcriptome. If supplied, *TopHat2* will map against the transcriptome first. Badly mapped and unmapped reads are aligned against the entire genome afterwards. The argument `addArgs` can be used to supply additional commandline arguments such as `--b2-very-sensitive`. See the *TopHat2* or *HISAT2* manual for more parameters.

```
topDir <- file.path(outDir, "mapping", "tophat")
map_tophat_res <- run_Tophat(
    files    = fq,
    index     = map_index,
    outDir    = topDir,
    is.paired = FALSE,
    cpus      = 2,
    workers   = 2
)
```

```
## TopHat2 - mapping ...
```

```
hiDir <- file.path(outDir, "mapping", "hisat")
map_hisat_res <- run_Hisat2(
    files     = fq,
    index     = map_index,
    outDir    = hiDir,
    is.paired = FALSE,
    cpus      = 2,
    workers   = 2
)
```

```
## HISAT2 - mapping ...
```

```
##
```

*MultiQC* (see next section) is able to process results from both mapping tools. Still, their reports should not be mixed. For this vignette and simplicity, the *TopHat2* results are removed.

```
unlink(topDir)
```

### 6 Gene Abundance Estimation

Gene abundance estimation refers to counting the number of reads mapping to pre-defined locations. In the context of transcriptomics, these location are usually genes or, more general, “features”. A feature is defined as an interval (start:stop) on a particular reference sequence. A gene composed of multiple exons may thus be defined by multiple features at distinct locations.

#### 6.1 Annotation

Genome annotations can be acquired in various ways. Read counting in *GEO2RNAseq* works best with files in GTF format. The Bioconductor package *rtracklayer* can be used to convert different kinds of gene annotation objects, e.g. *GRanges*, to GTF files. Each line in a GTF file defines a single feature. The annotation corresponding to the *Drosophila melanogaster* genome dm3 can be acquired and converted as follows:

```
library(TxDb.Dmelanogaster.UCSC.dm3.ensGene)
```

```
exbygene <- GenomicFeatures::exonsBy(TxDb.Dmelanogaster.UCSC.dm3.ensGene, "gene")
# convert GRangesList to GRanges object
exbygene <- unlist(exbygene)
rtracklayer::export(exbygene, file.path(outDir, "dm3.chr4.gtf"), format = "gtf")
```

An annotation file in GTF format is required for counting (see next section).

#### 6.2 Counting

Counting is the process of quantifying the number of reads assigned to each feature, e.g. genes, in the reference genome.

*GEO2RNAseq* uses *featureCounts* from the *Rsubread* package for counting. The annotation should be supplied in GTF format. Essential arguments are `gene_type` and `feature_type`. The `feature_type` refers to the 3rd column of the GTF file. This is usually “CDS”, “exon” or “gene”. By default, “exon” is used. The `gene_type` refers to the gene ID attribute in the rightmost column of the GTF file. It is “gene\_id” by default.

For example, two or more exons may be part of the same gene. In that case, all exons share the same ‘gene\_id’ attribute. In general, *featureCounts* will combine the counts of all ‘feature\_type’ entries with the same ‘gene\_type’ value. *Make sure to choose these two values correctly. Otherwise, each read mapping to two or more exons of the same gene may falsely be rejected as ambiguouse (because a read is only counted for one gene)*.

The GTF file produced in the example above uses “exon\_id” and “sequence\_feature”.

```
bamFiles <- map_hisat_res$files
gene_type <- "ID"
feature_type <- "sequence_feature"
count_res <- run_featureCounts(
    files       = bamFiles,
    annotation  = file.path(outDir, "dm3.chr4.gtf"),
    outDir      = file.path(outDir, "counting"),
    featureType = feature_type,
    IDtype      = gene_type,
    isPairedEnd = FALSE,
    cpus = 2,
    workers = 2
)
```

```
## featureCounts running... Paired: FALSE
```

```
count_res$summary
```

```
##                        Assigned Unassigned_Unmapped
## synthetic.trimo.fq.bam      335                   0
##                        Unassigned_MappingQuality Unassigned_Chimera
## synthetic.trimo.fq.bam                         0                  0
##                        Unassigned_FragmentLength Unassigned_Duplicate
## synthetic.trimo.fq.bam                         0                    0
##                        Unassigned_MultiMapping Unassigned_Secondary
## synthetic.trimo.fq.bam                     168                    0
##                        Unassigned_Nonjunction Unassigned_NoFeatures
## synthetic.trimo.fq.bam                      0                   571
##                        Unassigned_Overlapping_Length Unassigned_Ambiguity
## synthetic.trimo.fq.bam                             0                    3
```

```
head(count_res$counts[rowSums(count_res$counts) > 0,])
```

```
## FBgn0002521 FBgn0004607 FBgn0004859 FBgn0005558 FBgn0005561 FBgn0005666 
##           2          10           4           4           5          23
```

The function will also return a lot of annotation information. It contains, among other things, the length of genes (length of combined exons) as used by featureCounts.

```
gene_lengths <- count_res$anno$Length
counts <- count_res$counts
head(count_res$anno)
```

```
##        GeneID
## 1 FBgn0000003
## 2 FBgn0000008
## 3 FBgn0000014
## 4 FBgn0000015
## 5 FBgn0000017
## 6 FBgn0000018
##                                                                             Chr
## 1                                                                         chr3R
## 2 chr2R;chr2R;chr2R;chr2R;chr2R;chr2R;chr2R;chr2R;chr2R;chr2R;chr2R;chr2R;chr2R
## 3       chr3R;chr3R;chr3R;chr3R;chr3R;chr3R;chr3R;chr3R;chr3R;chr3R;chr3R;chr3R
## 4       chr3R;chr3R;chr3R;chr3R;chr3R;chr3R;chr3R;chr3R;chr3R;chr3R;chr3R;chr3R
## 5       chr3L;chr3L;chr3L;chr3L;chr3L;chr3L;chr3L;chr3L;chr3L;chr3L;chr3L;chr3L
## 6                                                                   chr2L;chr2L
##                                                                                                                  Start
## 1                                                                                                              2648220
## 2 18024494;18024496;18024938;18025505;18039159;18050410;18052282;18056749;18058283;18059587;18059821;18060002;18060002
## 3          12632936;12633349;12635854;12648144;12651773;12652857;12652857;12652857;12653461;12654643;12654643;12654643
## 4          12752932;12753395;12755360;12756838;12757124;12758093;12761649;12768975;12769504;12785694;12788881;12797481
## 5          16615470;16615470;16616737;16619373;16619628;16619843;16620022;16620602;16621274;16621763;16624477;16639910
## 6                                                                                                    10973443;10974268
##                                                                                                                    End
## 1                                                                                                              2648518
## 2 18024531;18024713;18025756;18025756;18039200;18051199;18052494;18058222;18059490;18059757;18059938;18060339;18060346
## 3          12635783;12635783;12636077;12648191;12652029;12653359;12653619;12653845;12653619;12655300;12655474;12655767
## 4          12755302;12755302;12755574;12757039;12757335;12760298;12761826;12769021;12769991;12786228;12789667;12797958
## 5          16616388;16618374;16618374;16619716;16619716;16619963;16620292;16621188;16621689;16622306;16624530;16640982
## 6                                                                                                    10974210;10975273
##                      Strand Length
## 1                         +    299
## 2 +;+;+;+;+;+;+;+;+;+;+;+;+   5400
## 3   -;-;-;-;-;-;-;-;-;-;-;-   5491
## 4   -;-;-;-;-;-;-;-;-;-;-;-   7719
## 5   -;-;-;-;-;-;-;-;-;-;-;-   6315
## 6                       -;-   1774
```

Usually, counts are normalized for further analysis. For example, TPM and RPKM values can be calculated on a matrix using the following functions:

```
tpm  <- get_tpm(counts, gene_lengths, colSums(counts))
rpkm <- get_rpkm(counts, gene_lengths, colSums(counts))
```

### 7 Mapping Statistics

Mapping statistics are, for example, how many raw input reads were available and how many were removed or remain unused after the processing steps until after counting. They also include an estimation of the genome coverage (the average number of reads assigned to each base of the total reference) and the exon coverage (the average number of reads assigned to each base of the exome).

Mapping statistics can be calculated with two different methods. Both methods differ only in the way they calculate the exon coverage.

The ‘estimate’ method uses the count matrix, genome length, and average read alignment length to estimate the exon coverage. This method is comparatively fast but requires a UNIX system. It reflects how many reads *are actually used* for the DEG Analysis. *featureCounts* does not use any read overlapping two or more genes. Statistical analysis is performed based on the counting table. Therefore, this calculation may reflect the exon coverage better with respect to later statistical analyses.

The ‘precise’ method determines the exact number of nucleotides mapped to exon regions. It uses several Bioconductor packages and requires sorted, indexed BAM files. Additionally, it does not consider results from *featureCounts*. Instead, only the information contained in the BAM files is used. This also means that reads mapping to multiple features are not excluded. The resulting exon coverage reflects how many reads *can potentially be used* for the DEG Analysis (if letting *featureCounts* also count such multi-overlapping reads).

```
mapping_stats_df <- calc_mapping_stats(
   bamFiles    = bamFiles,
   fqFiles     = system.file("extdata/synthetic.fastq.gz", package = "Geo2RNAseq"),
   anno        = file.path(outDir, "dm3.chr4.gtf"),
   numReads    = trim_res$input,
   numTrimmed  = trim_res$surviving,
   numNonrRNA  = NA,
   libSizes    = count_res$summary[,1],
   featureType = feature_type,
   paired      = FALSE,
   precise     = FALSE,
   remove.na   = FALSE,
   cpus = 2
)
mapping_stats_df
```

```
##                                                                                          fastq_files
## synthetic.fastq.gz /home/basti/.conda/envs/r-dev/lib/R/library/Geo2RNAseq/extdata/synthetic.fastq.gz
##                    reads_raw reads_trimmed percentage_reads_trimmed
## synthetic.fastq.gz      1000          1000                        0
##                    reads_non_rRNA percentage_non_rRNA_reads reads_mapped
## synthetic.fastq.gz             NA                        NA          993
##                    reads_unmapped percentage_reads_mapped
## synthetic.fastq.gz              7                    99.3
##                    percentage_total_read_loss genome_coverage
## synthetic.fastq.gz                        0.7      0.03672726
##                    reads_mapping_in_exons percentage_reads_mapping_in_exons
## synthetic.fastq.gz                    335                              33.5
##                    exon_coverage
## synthetic.fastq.gz  0.0005074782
```

The resulting object is a simpel data frame. It can be saved as simpel tsv or excel file like this:

```
write_count_table(
    file       = file.path(outDir, "mapping_stats"),
    counts     = mapping_stats_df,
    as.xls     = TRUE,
    rnames     = TRUE
)
```

#### 7.1 Explanation of Mapping Stats Table

The mapping statistics are made to give a brief overview over the mapping result. Each row represents a sample. The most important columns are ‘percentage reads mapped’, ‘percentage total read loss’ and ‘exon coverage’. ‘percentage reads mapped’ is just the fraction of the remaining input reads (after quality filtering and rRNA removal) that could be assigned to *at least on location*. A considerable amount of reads may map to multiple optimal locations (*multi mapping reads*). You should therefore check the MultiQC report in the next section (in our example used, the multi map rate is 8% for HISAT2). In fact, featureCounts will ignore such reads by default. This can therefore lead to a strong decline in exon coverage as well. Whether this is the case, or not, is also shown in the MultiQC report. ‘percentage total read loss’ calculates the number of reads lost until after mapping compared to the absolute number of raw input reads. High percentage values may indicate a fundamental problem with the sample reads (e.g. very bad quality on average or high rRNA content), the reference (wrong/bad reference genome) or filtering methods (quality cut-offs to strict, wrong parameter setup). ‘genome coverage’ and ‘exon coverage’ are very similar. These show how often, on average, a base in the reference is overlapped (“covered”) by reads. For genome coverage, the entire genome sequence and all mapped reads are considered. For exon coverage, only the exome size and only reads mapping into exon features are considered. A coverage of 1 means: the entire genome/exome is covered exactly once (on average). For any experiment dealing with gene expression analysis, exon coverage is the most important value. As mentioned before, the counting step may reduce another set of reads in addition to the mapping step. As a rule of thumb, one should aim for an exon coverage of at least 5-fold; around 10-fold and more would be perfect. Considerably more than 10-fold is not neccessary for gene expression analysis, but may be for other types of analyses.

### 8 MultiQC

MultiQC searches a directory and its subdirectories for various report files and combines them into a single interactive HTML report. For example, individual *FastQC* read quality charts are combined into a single chart for all samples. It will parse the reports and logs from *FastQC*, *Trimmomatic*, *HISAT2*, *TopHat2*, *SAMtools*, and *featureCounts*.

```
multiqc <- run_MultiQC(dir = outDir, config = get_MultiQC_config())
```

```
## See MultiQC report at '/home/basti/.conda/envs/r-dev/lib/R/library/Geo2RNAseq/extdata/outData/multiqc_report.html'
```

```
multiqc
```

```
## [1] "/home/basti/.conda/envs/r-dev/bin/multiqc --force -o '/home/basti/.conda/envs/r-dev/lib/R/library/Geo2RNAseq/extdata/outData' -e bowtie2  -c '/home/basti/.conda/envs/r-dev/lib/R/library/Geo2RNAseq/extdata/multiqc_config.yaml' '/home/basti/.conda/envs/r-dev/lib/R/library/Geo2RNAseq/extdata/outData'"
```

*MultiQC can integrate many more tools if the user chooses to add their own processing steps to the standard GEO2RNAseq pipeline*.

### 9 Clustering and PCA

#### 9.1 Load Count Data

*The following step is only done for this vignette. For your own data set, you may simply jump to the next section.*
In our example run, we have generated count data from raw FASTQ files. However, these previously generated count data is too small for the following analyses. Therefore, we load count data from the *airway* package.

In this experiment, four different cell lines of the smooth muscle from the human airway were treated with dexamethasone (DEX). DEX is a Glucocorticoid used for treating Asthma patients. The goal was to find differentially expressed genes in response to a treatment with DEX.

```
data(airway)
counts <- assay(airway) # FOR YOUR DATA, USE 'counts <- count_res$counts' !!!
gene_lengths <- width(IRanges::PartitioningByEnd(rowRanges(airway)))
# data info
metadata(rowRanges(airway))
```

```
## $genomeInfo
## $genomeInfo$`Db type`
## [1] "TranscriptDb"
## 
## $genomeInfo$`Supporting package`
## [1] "GenomicFeatures"
## 
## $genomeInfo$`Data source`
## [1] "BioMart"
## 
## $genomeInfo$Organism
## [1] "Homo sapiens"
## 
## $genomeInfo$`Resource URL`
## [1] "www.biomart.org:80"
## 
## $genomeInfo$`BioMart database`
## [1] "ensembl"
## 
## $genomeInfo$`BioMart database version`
## [1] "ENSEMBL GENES 75 (SANGER UK)"
## 
## $genomeInfo$`BioMart dataset`
## [1] "hsapiens_gene_ensembl"
## 
## $genomeInfo$`BioMart dataset description`
## [1] "Homo sapiens genes (GRCh37.p13)"
## 
## $genomeInfo$`BioMart dataset version`
## [1] "GRCh37.p13"
## 
## $genomeInfo$`Full dataset`
## [1] "yes"
## 
## $genomeInfo$`miRBase build ID`
## [1] NA
## 
## $genomeInfo$transcript_nrow
## [1] "215647"
## 
## $genomeInfo$exon_nrow
## [1] "745593"
## 
## $genomeInfo$cds_nrow
## [1] "537555"
## 
## $genomeInfo$`Db created by`
## [1] "GenomicFeatures package from Bioconductor"
## 
## $genomeInfo$`Creation time`
## [1] "2014-07-10 14:55:55 -0400 (Thu, 10 Jul 2014)"
## 
## $genomeInfo$`GenomicFeatures version at creation time`
## [1] "1.17.9"
## 
## $genomeInfo$`RSQLite version at creation time`
## [1] "0.11.4"
## 
## $genomeInfo$DBSCHEMAVERSION
## [1] "1.0"
```

A ‘dex’ and a ‘cell’ value is assigned to each sample. In the following, samples with ‘dex=trt’ will be considered replicates of one condition called ‘DEX’. Samples with ‘dex=untrt’ will be considered replicates of a second condition called ‘noDEX’.

#### 9.2 Hierarchical Clustering

Hierarchical clustering allows the user to group their data in a hierarchical dendrogram and to easily visualize groups within the dataset. This can reveal issues in the dataset, like potentially mislabeled samples, high or low variance within conditions, and batch effects. Two types of hierarchical clustering can be performed with *GEO2RNAseq*: (1) hierarchical clustering and (2) heat clustering, both based on raw count data or transformed/normalized data.

Transformations like MRN, CPM, RPKM, TPM or *user defined ones* can be used instead of raw count data. It only considers the data as it is without any assumptions. Additionally, variance stabilizing transformation (rlog or VST), as implemented in the *DESeq2* package, is available.

Clustering works best with a condition vector. It can be defined by the user through the `conds` argument or generated from the design matrix using `conditions_from_design()`. In case of the *airway* package, the ‘dex’ column can be used as condition vector. Sample names will be colored based on this vector, i.e. samples of the same condition will have the same color.

```
plotDir <- file.path(outDir, "result_plots")
conds  <- as.character(colData(airway)$dex)

# hierarchical clustering for MRN-normalized count values
# any kind of normalization may be applied to counts
clust_file <- file.path(plotDir, "hierarchical_clustering")
dist_matrix <- make_hclust_plot(
    clust_file,
    counts = counts,
    conds  = conds,
    norm   = "mrn"
)
dist_matrix
```

```
##            SRR1039508 SRR1039509 SRR1039512 SRR1039513 SRR1039516 SRR1039517
## SRR1039509   145.0890                                                       
## SRR1039512   138.0958   162.2127                                            
## SRR1039513   171.6175   157.7246   150.4109                                 
## SRR1039516   147.9897   169.2142   143.1500   176.1677                      
## SRR1039517   166.4550   155.9161   158.6804   156.2510   132.3423           
## SRR1039520   141.7821   168.5385   136.7716   170.4814   151.3985   168.0996
## SRR1039521   167.8245   152.3186   161.8590   146.8132   172.8556   152.7183
##            SRR1039520
## SRR1039509           
## SRR1039512           
## SRR1039513           
## SRR1039516           
## SRR1039517           
## SRR1039520           
## SRR1039521   148.4914
```

```
# heat clustering
heat_file <- file.path(plotDir, "heat_hierarchical_clustering")
dist_matrix <- make_heat_clustering_plot(
    heat_file,
    counts = counts,
    conds = conds,
    norm = "mrn"
)
dist_matrix
```

```
##            SRR1039508 SRR1039509 SRR1039512 SRR1039513 SRR1039516 SRR1039517
## SRR1039509   145.0890                                                       
## SRR1039512   138.0958   162.2127                                            
## SRR1039513   171.6175   157.7246   150.4109                                 
## SRR1039516   147.9897   169.2142   143.1500   176.1677                      
## SRR1039517   166.4550   155.9161   158.6804   156.2510   132.3423           
## SRR1039520   141.7821   168.5385   136.7716   170.4814   151.3985   168.0996
## SRR1039521   167.8245   152.3186   161.8590   146.8132   172.8556   152.7183
##            SRR1039520
## SRR1039509           
## SRR1039512           
## SRR1039513           
## SRR1039516           
## SRR1039517           
## SRR1039520           
## SRR1039521   148.4914
```

```
# the same plot with TPM
heat_file <- file.path(plotDir, "heat_hierarchical_clustering_tpm.pdf")
dist_matrix <- make_heat_clustering_plot(
    heat_file,
    counts = counts,
    conds = conds,
    norm = "tpm",
    geneLen = gene_lengths
)
dist_matrix
```

```
##            SRR1039508 SRR1039509 SRR1039512 SRR1039513 SRR1039516 SRR1039517
## SRR1039509   20727.17                                                       
## SRR1039512   31370.82   28739.52                                            
## SRR1039513   39031.48   28676.63   18935.69                                 
## SRR1039516   63734.95   60087.25   51410.86   55962.77                      
## SRR1039517   50387.10   44844.76   36323.39   38859.26   25140.67           
## SRR1039520   26338.42   20006.64   29262.33   33682.34   48329.99   37554.16
## SRR1039521   30253.31   23307.67   31349.63   30584.31   53318.71   38602.08
##            SRR1039520
## SRR1039509           
## SRR1039512           
## SRR1039513           
## SRR1039516           
## SRR1039517           
## SRR1039520           
## SRR1039521   17243.10
```

Hierarchical clustering of samples, based on MRN-normalized count values

Heat clustering of samples, based on MRN-normalized count values

#### 9.3 Correlation

Pairwise pearson correlation plots indicate how similar two samples are in regards to monotonicity. Ideally, replicates of the same condition should be very similar. If a design matrix is supplied, correlation is calculated between replicates of each possible condition, i.e. two plots per comparison. If only a count matrix is supplied, all pairwise sample correlations are determined. The identity is plotted as red line, the blue line indicates up- and down-expression. Instead of individual points, densities are plotted (roughly circular shapes). Data values are transformed by log_10_(x+1) to scale up low expression data.

For this vignette, only the first two samples are used.

```
corr <- make_correlation_plots(
    dat    = get_mrn(counts[,1:2]),
    outDir = file.path(plotDir, "COR"),
    prefix = "corr"
)
corr
```

```
## [[1]]
## [1] 0.988
```

Pairwise Pearson correlation of samples

Here, the resulting correlation for the two replicates of the treatment groups looks as follows: We can see that most data points are close to 0, but the highest expression of a gene after log transformation is around 6.

#### 9.4 Principle Component Analysis

The principle component analysis (PCA) is a dimension reduction method. In the context of RNA-seq data analysis, it is used to visualize the similarity (or non-similarity) of samples, based on the dataset’s variance. Like the hierarchical clustering, it may help to reveal issues in the dataset, too. *GEO2RNAseq* will only display the data based on the first two most informative axes, i.e. the two principal components (PCs), which explain most of the dataset’s variance. Because of the two dimensions, this PCA plot is also called “biplot”.

Ideally, replicates of the same condition should be dense and clearly separated from other conditions. “Misplaced” samples could be the indication for batch effects. For example, using the same sequencing flow cell for different conditions could result in adjacent samples in the PCA. Likewise, different measuring time points for the same condition could result in distant samples. To highlight this kind of special “treatment”, besides the actual conditions, a vector called ‘shapes’ can be use with the PCA. Different symbols are assigned to each group in ‘shapes’. Eclipses showing the variance of each group in 2d space can also be added. The ggplot object (‘g’) and the data used to create it (‘biplot’) is returned for further modification. Just check the contents of the ‘pca\_res’ list.

```
shapes <- as.character(colData(airway)$cell)
conds  <- as.character(colData(airway)$dex)
# Alternatively, this function can guess the conditions based on the design matrix
# conds <- conditions_from_design(design_matrix)

pca_file <- file.path(plotDir, "PCA.pdf")
pca_res <- make_PCA_plot(
    file   = pca_file,
    # designMatrix = design_matrix,
    counts = counts[, conds != "none"],
    conds  =  conds[  conds != "none"],
    shapes = shapes[  conds != "none"],
    norm = "tpm",
    add_eclipse = TRUE,
    geneLen = gene_lengths
)
```

Principal component analysis – PCA biplot

### 10 Design Matrix

The design matrix is used to describe pairwise comparisons for the analysis of differentially expressed genes (DEGs), see next section.

In essence, one only needs to define which samples are in the *treatment* and in the *control* group. Each column in the matrix represents one comparison and each row represents a sample. To make it compatible with count data, the row names of the design matrix should be identical to the column names of the count matrix: `rownames(design) == colnames(counts)`. Each column name should have the form “<A>\_VS\_<B>”, where <A> and <B> are variable parts. <A> is always considered *treatment* and <B> *control*. The two conditions will be compared to identify significantly differentially expressed genes. The column names are also used to name the output files.

The matrix can be created by hand or be calculated based on common comparisons. The latter can be done using a function called ‘createDesignMatrix’. It would be best to check out the example for this Funktion to see how it may be used. In brief, it uses a metadata data.frame to enumerate comparisons. It uses either a specified column describing condtions, or time points, or a combination of both. The latter can create an extreme amount of comparisons very fast. Therefore, grouping of values and different modes are available. The following code will create the design matrix for the airway dataset.

```
createDesignMatrix(colData(airway), condCol = "dex")
```

```
## $dm
##           trt_VS_untrt
## SRS508568 "control"   
## SRS508567 "treatment" 
## SRS508571 "control"   
## SRS508572 "treatment" 
## SRS508575 "control"   
## SRS508576 "treatment" 
## SRS508579 "control"   
## SRS508580 "treatment" 
## 
## $tests
## [1] "trt_VS_untrt"
## 
## $meta
## DataFrame with 8 rows and 9 columns
##            SampleName     cell         dex    albut        Run avgLength
##              <factor> <factor> <character> <factor>   <factor> <integer>
## SRR1039508 GSM1275862   N61311       untrt    untrt SRR1039508       126
## SRR1039509 GSM1275863   N61311         trt    untrt SRR1039509       126
## SRR1039512 GSM1275866  N052611       untrt    untrt SRR1039512       126
## SRR1039513 GSM1275867  N052611         trt    untrt SRR1039513        87
## SRR1039516 GSM1275870  N080611       untrt    untrt SRR1039516       120
## SRR1039517 GSM1275871  N080611         trt    untrt SRR1039517       126
## SRR1039520 GSM1275874  N061011       untrt    untrt SRR1039520       101
## SRR1039521 GSM1275875  N061011         trt    untrt SRR1039521        98
##            Experiment    Sample    BioSample
##              <factor>  <factor>     <factor>
## SRR1039508  SRX384345 SRS508568 SAMN02422669
## SRR1039509  SRX384346 SRS508567 SAMN02422675
## SRR1039512  SRX384349 SRS508571 SAMN02422678
## SRR1039513  SRX384350 SRS508572 SAMN02422670
## SRR1039516  SRX384353 SRS508575 SAMN02422682
## SRR1039517  SRX384354 SRS508576 SAMN02422673
## SRR1039520  SRX384357 SRS508579 SAMN02422683
## SRR1039521  SRX384358 SRS508580 SAMN02422677
```

If not already defined in the metadata or created automatically by ‘createDesignMatrix’, the following code creates a new design matrix “manually”.

```
# design matrix
design_matrix <- matrix("none", ncol = 1, nrow = ncol(counts))
colnames(design_matrix) <- "DEX_VS_noDEX"
rownames(design_matrix) <- colnames(counts)
design_matrix[colData(airway)$dex == "trt",   1] <- "treatment"
design_matrix[colData(airway)$dex == "untrt", 1] <- "control"
design_matrix
```

```
##            DEX_VS_noDEX
## SRR1039508 "control"   
## SRR1039509 "treatment" 
## SRR1039512 "control"   
## SRR1039513 "treatment" 
## SRR1039516 "control"   
## SRR1039517 "treatment" 
## SRR1039520 "control"   
## SRR1039521 "treatment"
```

The condition vector used by the clustering and PCA methods can be generated from the design matrix using `conditions_from_design()`. Be aware that this function is experimental and may give unexpected results for arbitrary sample names. The design matrix produces the following condition vector.

```
conditions_from_design(design_matrix)
```

```
## [1] "noDEX" "DEX"   "noDEX" "DEX"   "noDEX" "DEX"   "noDEX" "DEX"
```

### 11 DEG Analysis

The analysis of differentially expressed genes (DEGs) can be performed with a variety of tools. By default, *DESeq*, *DESeq2*, *edgeR* and *limma* are used. Additionally, *NOISeq*, *baySeq*, *PoissonSeq* and SAMseq (from the *samr* package) are available.

The p-value cut-off is set to 0.01 by default. Optionally, a log_2_ fold change cut-off can be set. If defined, genes must pass both cut-offs to be considered significant. The cut-offs are also integrated into the resulting volcano plot(s). A volcano plot displays p-values versus log_2_ fold changes and gives a brief DEG overview for the compared conditions.

In the following, *DESeq* and *DESeq2* are excluded to reduce computation time.

```
degDir <- file.path(outDir, "diff_exp_genes")
tools <- c("edgeR", "limma")
deg_res <- calculate_DEGs(
    counts       = counts,
    geneLengths  = gene_lengths,
    libSizes     = colSums(counts),
    designMatrix = design_matrix,
    pValCut      = 0.01,
    logfcCut     = NA,
    tools        = tools,
    outDir       = degDir,
    cpus         = 2,
    workers      = 2
)
```

```
head(deg_res$DEGs$DEX_VS_noDEX)
```

```
##                              id mean_A_mrn  mean_B_mrn log2_fc_mrn
## ENSG00000000003 ENSG00000000003  616.69697  802.507368 -0.37995288
## ENSG00000000419 ENSG00000000419  558.08806  484.507740  0.20397307
## ENSG00000000457 ENSG00000000457  240.81196  235.514117  0.03209349
## ENSG00000000460 ENSG00000000460   56.54071   61.324557 -0.11717492
## ENSG00000000938 ENSG00000000938    1.00000    1.636197 -0.71034625
## ENSG00000000971 ENSG00000000971 6700.38620 4936.319535  0.44080847
##                 mean_A_tpm mean_B_tpm log2_fc_tpm mean_A_rpkm mean_B_rpkm
## ENSG00000000003  16.304698  21.804065 -0.41930938  1644.01346 2267.856462
## ENSG00000000419   9.143781   8.385502  0.12489361   874.96554  802.197355
## ENSG00000000457   4.386233   4.493572 -0.03488014   364.28206  377.473499
## ENSG00000000460   1.328269   1.374163 -0.04900586    36.10929   41.448836
## ENSG00000000938   1.000000   1.010316 -0.01480599     1.00000    2.151944
## ENSG00000000971  63.978258  49.294057  0.37616797  6731.96803 5260.647013
##                   fc_rpkm log2_fc_rpkm Limma Limma_adj_pval EdgeR
## ENSG00000000003 0.7249195  -0.46410722 FALSE      0.2396723 FALSE
## ENSG00000000419 1.0907111   0.12526899 FALSE      0.3939658 FALSE
## ENSG00000000457 0.9650533  -0.05131942 FALSE      0.9231674 FALSE
## ENSG00000000460 0.8711774  -0.19896160 FALSE      0.9081196 FALSE
## ENSG00000000938 0.4646962  -1.10564022 FALSE      0.5572630 FALSE
## ENSG00000000971 1.2796844   0.35578808 FALSE      0.4670917 FALSE
##                 EdgeR_adj_pval
## ENSG00000000003      0.4358943
## ENSG00000000419      1.0000000
## ENSG00000000457      1.0000000
## ENSG00000000460      1.0000000
## ENSG00000000938      0.9841073
## ENSG00000000971      0.5612470
```

Volcano plot

Different DEG tools often give different results, i.e. different sets of DEGs according to the assigned p-values. If using more than one tool, `calculate_DEGs()` also creates a Venn diagram and an intersection bar plot for each comparison. With more and more tools to compare, Venn diagrams become rather ugly and eventually almost impossible to draw. The intersection bar plots are an easy-to-read alternative if comparing more than four tools.

Venn diagram for different DEG tools

Intersection bar plot for different DEG tools

The log_2_ fold changes in the tables created by `calculate_DEGs()` are calculated by transforming (normalizing) the raw count data into MRN (default), TPM or RPKM values and calculating the mean of replicates. The log_2_ fold change for gene *i* is defined as:

*log_2_ ( treatmeant(i) / control(i) )*

When many comparisons were conducted, you may like to see the distribution of DEGs for each test. We present this in form of a bar chart, where the number of up- and down-regulated as well as the total number is displayed. Such a plot can be created with the function `make_deg_overview_plot`.

```
deg_overview <- make_deg_overview_plot(
    degs  = deg_res$DEGs,
    tools = tools
)
```

DEGs Overview plot

### 12 Updating Metadata

The metadata is stored as a simple `data.frame`. Updating this `data.frame` can be done in three ways: (1) write it to disk and update it using tools like Excel or LibreOffice Calc, (2) modify it like any other `data.frame` or (3) use the `updateMeta` function from *GEO2RNAseq*. The latter one will be demonstrated based on the results of this vignette.

For this, the metadata should be loaded using a parser function (functions starting with “parse”). Here, we load a metadata table in SDRF format:

```
SDRFinfo <- parse_SDRF(file.path(pkgDir, "GEO", "SDRF_template.tsv"))
```

```
## Fold Change Assignment column not found. Ignored.
```

We performed quality trimming, mapping and read counting on the data. Differential gene expression analysis was peformed on a different dataset, but will also be added for demonstration.

```
SDRFinfo <- updateMeta(
    metaRes = SDRFinfo,
    toolRes = fq_raw_res
)
```

```
## Selected rank: '0'
```

```
SDRFinfo <- updateMeta(
    metaRes = SDRFinfo,
    toolRes = trim_res,
    rank    = 2
)
SDRFinfo <- updateMeta(
    metaRes = SDRFinfo,
    toolRes = map_hisat_res,
    rank    = 3
)
SDRFinfo <- updateMeta(
    metaRes = SDRFinfo,
    toolRes = count_res,
    rank    = 4
)
SDRFinfo <- updateMeta(
    metaRes = SDRFinfo,
    toolRes = deg_res,
    rank    = 5
)
```

The ‘rank’ argument is optional, but should be set. The newly added entries can be found in the rightmost columns of the resulting matrix.

As before, you may write the changes to the metadata table down to hard drive using:

```
meta <- write_meta_table(SDRFinfo$table, as.xls = FALSE, outDir = outDir, overwrite = T)
```

```
## Metadata files already exist. Overwrite enabled...
```

```
meta
```

```
## [1] "/home/basti/.conda/envs/r-dev/lib/R/library/Geo2RNAseq/extdata/outData/meta_data.csv"
```

### 13 Archive Results

It is advised to save the R workspace, i.e. the value of all variables (R objects), to disk.

```
save.image(file.path(outDir, "R_workspace.RData")) # save workspace
```

One can do this, for example, after each pre-processing step. For archiving purposes, one should do it at least once after finishing the pipeline and keep the file, together with the RNA-seq raw data and main analysis results, at a save place.

### 14 Example Pre-processing Workflows (Pipelines)

The workflow presented in this vignette can be used as an example RNA-seq pipeline/workflow. More comprehensive and highly recommended pipeline scripts, which utilize almost all functions of *GEO2RNAseq*, are shipped with the package. Feel free to modify them to fit your needs and datasets!

```
system.file("extdata/pipelines", package = "Geo2RNAseq")
```

```
## [1] "/home/basti/.conda/envs/r-dev/lib/R/library/Geo2RNAseq/extdata/pipelines"
```

### 15 Session Info

```
sessionInfo()
```

```
## R version 3.4.1 (2017-06-30)
## Platform: x86_64-pc-linux-gnu (64-bit)
## Running under: Ubuntu 18.04.2 LTS
## 
## Matrix products: default
## BLAS: /home/basti/.conda/envs/r-dev/lib/R/lib/libRblas.so
## LAPACK: /home/basti/.conda/envs/r-dev/lib/R/lib/libRlapack.so
## 
## locale:
##  [1] LC_CTYPE=en_US.UTF-8       LC_NUMERIC=C              
##  [3] LC_TIME=de_DE.UTF-8        LC_COLLATE=en_US.UTF-8    
##  [5] LC_MONETARY=de_DE.UTF-8    LC_MESSAGES=en_US.UTF-8   
##  [7] LC_PAPER=de_DE.UTF-8       LC_NAME=C                 
##  [9] LC_ADDRESS=C               LC_TELEPHONE=C            
## [11] LC_MEASUREMENT=de_DE.UTF-8 LC_IDENTIFICATION=C       
## 
## attached base packages:
##  [1] parallel  stats4    grid      stats     graphics  grDevices utils    
##  [8] datasets  methods   base     
## 
## other attached packages:
##  [1] airway_0.112.0                           
##  [2] SummarizedExperiment_1.8.0               
##  [3] DelayedArray_0.4.1                       
##  [4] matrixStats_0.52.2                       
##  [5] TxDb.Dmelanogaster.UCSC.dm3.ensGene_3.2.2
##  [6] GenomicFeatures_1.30.3                   
##  [7] AnnotationDbi_1.40.0                     
##  [8] Biobase_2.38.0                           
##  [9] GenomicRanges_1.30.0                     
## [10] GenomeInfoDb_1.14.0                      
## [11] IRanges_2.12.0                           
## [12] knitr_1.16                               
## [13] Geo2RNAseq_0.99.13                       
## [14] S4Vectors_0.16.0                         
## [15] BiocGenerics_0.24.0                      
## [16] BiocStyle_2.6.0                          
## 
## loaded via a namespace (and not attached):
##   [1] backports_1.1.0          Hmisc_4.0-3             
##   [3] plyr_1.8.4               lazyeval_0.2.0          
##   [5] splines_3.4.1            crosstalk_1.0.0         
##   [7] BiocParallel_1.12.0      NOISeq_2.22.0           
##   [9] ggplot2_2.2.1            digest_0.6.12           
##  [11] BiocInstaller_1.28.0     htmltools_0.3.6         
##  [13] viridis_0.4.0            gdata_2.18.0            
##  [15] magrittr_1.5             checkmate_1.8.2         
##  [17] memoise_1.1.0            cluster_2.0.6           
##  [19] limma_3.34.6             Biostrings_2.46.0       
##  [21] annotate_1.56.0          R.utils_2.5.0           
##  [23] prettyunits_1.0.2        colorspace_1.3-2        
##  [25] blob_1.1.0               WriteXLS_4.0.0          
##  [27] dplyr_0.7.0              xfun_0.1                
##  [29] jsonlite_1.5             RCurl_1.95-4.8          
##  [31] genefilter_1.60.0        glue_1.1.1              
##  [33] survival_2.41-3          gtable_0.2.0            
##  [35] zlibbioc_1.24.0          XVector_0.18.0          
##  [37] Rsubread_1.28.0          UpSetR_1.3.3            
##  [39] kernlab_0.9-25           prabclus_2.2-6          
##  [41] DEoptimR_1.0-8           abind_1.4-5             
##  [43] scales_0.4.1             DESeq_1.30.0            
##  [45] futile.options_1.0.0     pheatmap_1.0.8          
##  [47] mvtnorm_1.0-6            DBI_0.6-1               
##  [49] edgeR_3.20.7             Rcpp_0.12.11            
##  [51] viridisLite_0.2.0        xtable_1.8-2            
##  [53] progress_1.1.2           htmlTable_1.9           
##  [55] foreign_0.8-68           bit_1.1-12              
##  [57] mclust_5.4               preprocessCore_1.40.0   
##  [59] Formula_1.2-1            baySeq_2.12.0           
##  [61] htmlwidgets_1.0          httr_1.2.1              
##  [63] RColorBrewer_1.1-2       fpc_2.1-10              
##  [65] acepack_1.4.1            modeltools_0.2-21       
##  [67] reshape_0.8.6            pkgconfig_2.0.1         
##  [69] XML_3.98-1.7             R.methodsS3_1.7.1       
##  [71] flexmix_2.3-14           nnet_7.3-12             
##  [73] locfit_1.5-9.1           labeling_0.3            
##  [75] rlang_0.1.1              munsell_0.4.3           
##  [77] tools_3.4.1              RSQLite_2.0             
##  [79] devtools_1.13.2          evaluate_0.10.1         
##  [81] stringr_1.2.0            yaml_2.1.14             
##  [83] bit64_0.9-7              robustbase_0.92-7       
##  [85] purrr_0.2.2              dendextend_1.5.2        
##  [87] nlme_3.1-131             mime_0.5                
##  [89] whisker_0.3-2            R.oo_1.21.0             
##  [91] biomaRt_2.34.2           compiler_3.4.1          
##  [93] plotly_4.7.1             affyio_1.48.0           
##  [95] tibble_1.3.3             geneplotter_1.56.0      
##  [97] stringi_1.1.6            highr_0.6               
##  [99] futile.logger_1.4.3      lattice_0.20-35         
## [101] trimcluster_0.1-2        Matrix_1.2-10           
## [103] samr_2.0                 data.table_1.10.4       
## [105] bitops_1.0-6             httpuv_1.3.3            
## [107] rtracklayer_1.38.0       affy_1.56.0             
## [109] R6_2.2.1                 latticeExtra_0.6-28     
## [111] hwriter_1.3.2            bookdown_0.6            
## [113] RMySQL_0.10.13           ShortRead_1.36.0        
## [115] KernSmooth_2.23-15       gridExtra_2.2.1         
## [117] lambda.r_1.2             MASS_7.3-47             
## [119] gtools_3.5.0             assertthat_0.2.0        
## [121] DESeq2_1.18.1            rprojroot_1.2           
## [123] withr_1.0.2              GenomicAlignments_1.14.0
## [125] Rsamtools_1.30.0         GenomeInfoDbData_1.0.0  
## [127] diptest_0.75-7           mgcv_1.8-17             
## [129] VennDiagram_1.6.18       rpart_4.1-11            
## [131] tidyr_0.6.3              class_7.3-14            
## [133] rmarkdown_1.8            shiny_1.0.3             
## [135] base64enc_0.1-3
```
