## Supplementary material for "GEO2RNAseq: An easy-to-use R pipeline for complete pre-processing of RNA-seq data": Minimal Pipeline Example

#!/usr/bin/env R

# This could should be executed from within the conda environment 'geo2rnaseq'
# The environment can be activated in any console by calling:
#   'source activate geo2RNAseq'
#
# This pipeline is fully modular and can be started at any given point
# Mostly the paths to input files (e.g. FASTQ, BAM, ...) are required

library("Geo2RNAseq")

### This part is only relevant if datasets from GEO are being processed

# Download data & parse meta data
geo_dat <- getGEOdata(accession = "GSE55663", outDir = "GSE55663/")
outDir <- "."

# Grab the SDRF formatted data.frame
# We complement this table semi-automatically with information of the processing done in this script
SDRF <- geo_dat$SDRF

# Conditions can be read from this SDRF table
head(SDRF$Extract.Name)

# For later visualization, we give these groups more readable names
conds <- c(rep("untreated", 3), rep("caspofungin", 3), rep("siamycin", 3), rep("CAS+SIA", 3))

# We check if we have single-end, paired-end, or both kinds of FASTQ files
head(geo_dat$se_files)
head(geo_dat$pe_files)


### Here, we start with FASTQ file processing


# Only single-end files were parsed, so we process all data in single-end mode from here on
paired <- F
fastq_files <- geo_dat$se_files # paths to downloaded FASTQ files

# Quality check of raw reads
fq_raw_res <- run_FastQC(
    files  = fastq_files,
    outDir = file.path("quality", "raw")
)

## Quality and adapter trimming
# Default Illumina adapters are part of Trimmomatic and are shipped with this package
# Additionally, we want our trimmed output files to be gz compressed
# Some tools require uncompressed FASTQ files, so we leave this optional
trim_res <- run_Trimmomatic(
    files     = fastq_files,
    outDir    = file.path("fastq"),
    is.paired = paired,
    compress  = TRUE
)

trim_res # contains executed commands, number of input reads, number of output reads and paths to trimmed FASTQ files
fastq_files <- trim_res$files # we continue with trimmed files


# Quality check of trimmed reads
fq_raw_res <- run_FastQC(
    files  = fastq_files,
    outDir = file.path("quality", "filter"),
    extend = TRUE
)

# Alignment of reads against appropriate genome reference
# We downloaded the genome reference of Aspergillus fumigatus from http://aspergillusgenome.org/
# Genome index was created using 'make_HiSat2_index()'
ref_genome <- "A_fumigatus_A1163_05_03_2018/genome/Aspergillus_fumigatusa1163.CADRE.dna.nonchromosomal.fa"
map_index  <- "A_fumigatus_A1163_05_03_2018/index_hisat2/Aspergillus_fumigatusa1163.CADRE.dna.nonchromosomal"
anno       <- "A_fumigatus_A1163_05_03_2018/annotation/Aspergillus_fumigatusa1163.CADRE.38.gtf"

map_hisat_res <- run_Hisat2(
    files     = fastq_files,
    index     = map_index,
    outDir    = file.path("mapping", "hisat"),
    is.paired = paired
)
bamFiles <- map_hisat_res$files # we continue with mapping files in BAM format


# Gene abundance estimation by read counting
# 'gene_type' and 'feature_type' must be chosen based on the annotation GTF file
# Here, we want reads covering multiple exons of the same gene to be summed up at gene level
count_res <- run_featureCounts(
    files       = bamFiles,
    annotation  = anno,
    outDir      = file.path("counting"),
    featureType = "exon",
    IDtype      = "gene_id",
    isPairedEnd = paired
)

# We should investigate briefly if the mapping was successful
# Comprehensive mapping statistics are created next
count_res$summary

# Get lengths of genes (number of nucleotides)
# This information is often required for normalization methods (e.g. RPKM)
gene_lengths <- count_res$anno$Length
# Get counting table, i.e. the number of reads mapped per gene per sample
counts <- count_res$counts
# Get effective library sizes, i.e. the total number of reads mapping to exons per samples
libSizes <- count_res$summary[,1]
# Instead of just samples names, we give the samples more informative names
# We know from the meta data SDRF that each condition contains 4 replicates
# Samples were ordered by condition first, then replicates
colnames(counts) <- paste0(conds, "_", paste0("rep", rep(1:3, 4)))

# Calculate mapping stats
# The more information we have, the more complete is the output table
# We feed in all information that we have
mapping_stats_df <- calc_mapping_stats(
   bamFiles    = bamFiles,
   fqFiles     = geo_dat$se_files,
   anno        = anno,
   numReads    = trim_res$input,
   numTrimmed  = trim_res$surviving,
   numNonrRNA  = NA,
   libSizes    = libSizes,
   featureType = "exon",
   paired      = paired,
   precise     = FALSE
)

# Save this table
# This table should be investigates first before proceeding
write_count_table(
    file   = file.path(outDir, "mapping_stats"),
    counts = mapping_stats_df,
    as.xls = TRUE,
    rnames = TRUE
)


# MultiQC combines all output logs and other information and creates an interactive HTML site
run_MultiQC(dir = outDir, config = get_MultiQC_config());

# These steps are fully optional
# We want to create the most basic analytical plots
plotDir <- file.path("result_plots")

# Creates a simple dendrogram with colored labels
# Coloring is based on the given 'conds' variable
# uses "Median-by-Ratio normalization" by default
make_hclust_plot(
    file   = file.path(plotDir, "hierarchical_clustering_counts"),
    main   = "Hierarchical Clustering MRN - UPGMA",
    counts = counts,
    conds  = conds
)


# Creates a heatmap with dendrograms on the borders
# uses "Median-by-Ratio normalization" by default
make_heat_clustering_plot(
    file   = file.path(plotDir, "heat_hierarchical_clustering_counts.mrn.pdf"),
    main   = "Hierarchical Clustering Counts",
    counts = counts,
    conds  = conds
)


# Principle Component Analysis
# These plots are often used to find out which samples are more similar to one another
# We often also want to see how diverse replicates of each condition are
# Therefore, we plot replicates using different symbols ('shapes') and conditions using different colors
# uses "Median-by-Ratio normalization" by default
shapes <- rep(1:4, each = 3)
pca_res <- make_PCA_plot(
    file        = file.path("PCA.pdf"), # output file names
    counts      = counts,
    conds       =  conds,
    shapes      = shapes,
    add_eclipse = TRUE    # plot 95% confidence intervals for groups defined in 'conds'
)


# Create design matrix
# We can auto-generate pairwise tests between conditions and different sample time points
# Here, we only have treatment conditions
# We add our simplified conditions to the meta data SDRF data.frame first
SDRF$condition <- conds
# the function detects potential naming issues and fixes them automatically
# if that happens, design$meta may be added to our SDRF object
design <- createDesignMatrix(SDRF, condCol = "condition")
# This is not the case here, so we just extract the actual contrast matrix
design_matrix <- design$dm
# The samples in the design matrix (rows) must have the same names as in the counting matrix
rownames(design_matrix) <- colnames(counts)
# We also add these to our meta data
SDRF <- cbind(SDRF, design_matrix)

# Calculate differentially expressed genes
# Every column of the design matrix is handled as independent test
# This function also creates some plots (e.g. volcano, intersection bar)
deg_res <- calculate_DEGs(
    counts       = counts,
    geneLengths  = gene_lengths, # we describe gene abundances in RPKM, TPM and MRN, so we need gene length information
    libSizes     = libSizes,
    designMatrix = design_matrix,
    pValCut      = 0.01,
    logfcCut     = 1,
    outDir       = file.path("diff_exp_genes"),
    cpus         = 2,
    workers      = 2
)

# Create a summary plot that simply displays how many DEGs were found
deg_overview <- make_deg_overview_plot(
    degs  = deg_res$DEGs,
    tools = c("DESeq", "DESeq2", "limma", "edgeR")
)

# At the end, we update the meta data
# Since the SDRF is a data.frame, we can add any amount of information here
# But at least, we shall add the information of the tools, the commands, and so on, to this table
SDRFinfo <- geo_dat
SDRFinfo$SDRF <- SDRF

SDRFinfo <- updateMeta(
    metaRes = SDRFinfo,
    toolRes = fq_raw_res,
    rank = 0
)
SDRFinfo <- updateMeta(
    metaRes = SDRFinfo,
    toolRes = trim_res,
    rank    = 1
)
SDRFinfo <- updateMeta(
    metaRes = SDRFinfo,
    toolRes = map_hisat_res,
    rank    = 2
)
SDRFinfo <- updateMeta(
    metaRes = SDRFinfo,
    toolRes = count_res,
    rank    = 3
)
SDRFinfo <- updateMeta(
    metaRes = SDRFinfo,
    toolRes = deg_res,
    rank    = 4
)

# save updated meta data to hard drive
write_meta_table(
    SDRFinfo$SDRF,
    as.xls = FALSE,
    outDir = outDir,
    overwrite = T
)

# save final workspace
save.image(file.path(outDir, "R_workspace.RData"))
